# Microdosing ketamine in *Drosophila* does not inhibit SERT like SSRIs, but causes behavioral changes mediated by glutamate and serotonin receptors

**DOI:** 10.1101/2023.11.07.566121

**Authors:** Kelly E. Dunham, Kani H. Khaled, Leah Weizman, B. Jill Venton

## Abstract

Recently, the FDA approved microdosing ketamine for treatment resistant depression. Traditional antidepressants, like serotonin selective reuptake inhibitors (SSRIs), block serotonin reuptake, but it is not clear if ketamine blocks serotonin reuptake. Here, we tested the effects of feeding ketamine and SSRIs to *Drosophila melanogaster* larvae, which has a similar serotonin system to mammals, and is a good model to track depression behaviors, such as locomotion and feeding. Fast-scan cyclic voltammetry (FSCV) was used to measure optogenetically-stimulated serotonin changes, and locomotion tracking software and blue dye feeding to monitor behavior. We fed larvae various doses (1-100 mM) of antidepressants for 24 hours and found that 1 mM ketamine did not affect serotonin, but increased locomotion and feeding. Low doses (≤ 10 mM) of escitalopram and fluoxetine inhibited dSERT and also increased feeding and locomotion behaviors. At 100 mM, ketamine inhibited dSERT and increased serotonin concentrations, but decreased locomotion and feeding due to its anesthetic properties. Since microdosing ketamine causes behavioral effects, we also investigated behavior changes with low doses of other NMDA receptor antagonists and 5-HT_1A and 2_ agonists, which are other possible sites for ketamine action. NMDA receptor antagonism increased feeding, while serotonin receptor agonism increased locomotion, which could explain these effects with ketamine. Ultimately, this work shows that *Drosophila* is a good model to discern antidepressant mechanisms, and that ketamine does not work on dSERT like SSRIs at microdoses, but affects behavior with other mechanisms.

**Graphical abstract:** 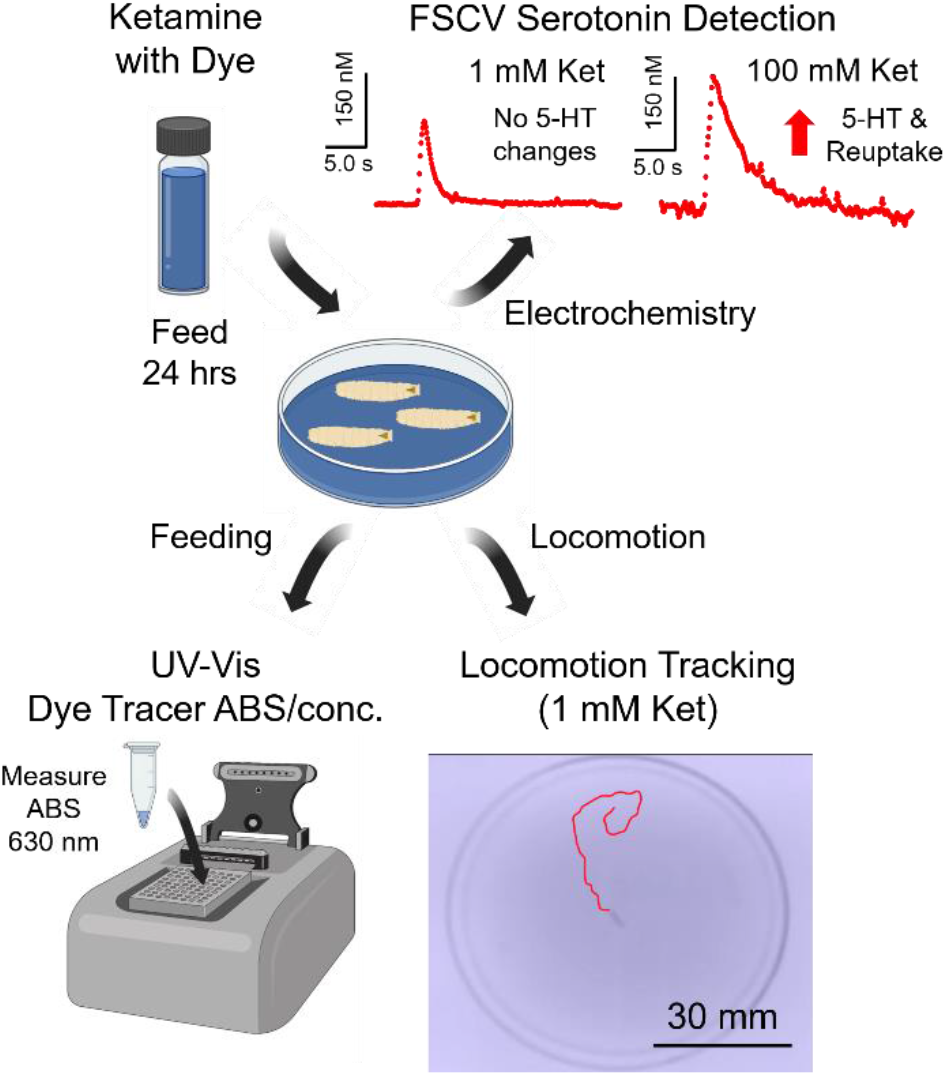

Microdosing ketamine is a novel depression treatment, but it is not clear how it changes serotonin in real-time. *Drosophila melanogaster* (fruit fly) is a good model to study depression behaviors. Here, we used fast-scan cyclic voltammetry, video tracking, and feeding assays to measure serotonin and behavior after feeding ketamine and SSRIs to larvae. At microdoses, ketamine did not affect serotonin, which was different from SSRIs. However, higher doses inhibited dSERT. Locomotion and feeding changes were also dose-dependent, and we saw separate effects with NMDA and serotonin receptor drugs. This work facilitates future behavioral and pharmacological testing with ketamine using *Drosophila*.

## Introduction

In 2019, the FDA approved microdosing ketamine to treat major depressive disorder (MDD) and treatment resistant depression (TRD) (Berman et al., 2000; Kraus et al., 2019; Ruberto et al., 2020). Patients with these diagnoses often fail to positively respond to at least 2 antidepressants, which are commonly selective serotonin reuptake inhibitors (SSRIs). Commonly, ketamine is used as an anesthetic at higher doses for mammals in surgeries, but sub-anesthetic doses produce psychedelic and dissociative effects (Kraus et al., 2019). For microdose treatments, patients are intravenously given low doses (0.5-1.0 mg/kg) over a period of several hours (Kraus et al., 2019). Interestingly, rapid-onset positive antidepressant effects have been reported within 4 hours that persist for up to 1-2 weeks (Kraus et al., 2019), which is remarkably different than conventional SSRIs that take many weeks to see initial improvements. Formally, ketamine is classified as a noncompetitive NMDA antagonist that blocks glutamate receptors (Kraus et al., 2019; Li, 2020). However, it also activates opioid receptors (Kraus et al., 2019; West et al., 2023), and binds to a variety of receptors and transporters in the brain, including serotonin (5-hydroxytryptophan, 5-HT) and dopamine (Bowman et al., 2020; Kraus et al., 2019; Li, 2020). Ketamine activates serotonin receptors, especially the 5-HT_1_ and 5-HT _2_ subtypes, which increase serotonin concentrations in the brain (Spies et al., 2018). However, it is not clear if ketamine blocks serotonin reuptake like an SSRI (Dunham & Venton, 2022; Owens et al., 2001; Ribaudo et al., 2021; Wood et al., 2014), since several studies using PET imaging and binding affinity assays have shown differing results (Barann et al., 2014; Can et al., 2016; Pham et al., 2017; Spies et al., 2018). Furthermore, it is not well understood if microdosing ketamine produces behavior changes that are due to activation of serotonin or other neurotransmitter systems (Hidalgo et al., 2017; Majeed et al., 2016; Silva et al., 2014).

*Drosophila melanogaster* is an ideal model organism to elucidate serotonin and behavior changes with antidepressants because their brain regions have been extensively characterized and they have homologous serotonin systems to mammals, including the serotonin transporter (SERT) and serotonin receptors (Dunham & Venton, 2022; Johnson et al., 2009; Kasture et al., 2018). A recent study showed that high, anesthetic ketamine doses are genotoxic to flies (Koksal & Gürbüzel, 2015), but sub-anesthetic doses have not been widely explored. There is an exquisite genetic toolbox available to modify the *Drosophila* genome to interrogate the effects of genetic modifications or knockdown of genes (Hales et al., 2015; Jeibmann & Paulus, 2009; Lawal et al., 2014; Roberts, 2006). In addition, *Drosophila* is a good model system for pharmacology screenings to understand antidepressant and anesthetic responses (Lawal et al., 2014; Martin & Krantz, 2014). *Drosophila* are also capable of performing complex behaviors that can be studied with simple assays after feeding antidepressants (Kasture et al., 2018). For example, climbing in adults is depressed by genetic manipulations or uncontrollable mechanical stress, and this depressive behavior can be alleviated with SSRIs, like fluoxetine (Kasture et al., 2018; Ries et al., 2017). Video tracking software can also be used to compare changes with different drugs or genetic mutations (Dumitrescu et al., 2023; Lawal et al., 2014; Majeed et al., 2016; Ries et al., 2017; Silva et al., 2014), including crawling in larvae (Ries et al., 2017; Tadres & Louis, 2020). For example, the Louis lab created the Raspberry Pi virtual reality system (PiVR) that monitors the movements of *Drosophila* larvae in an arena with video tracking equipment (Tadres & Louis, 2020). Several analytical behavior assays have also been created to measure feeding in *Drosophila* (Hidalgo et al., 2017; Majeed et al., 2016; Shell et al., 2018; Silva et al., 2014). Specifically, the Grotewiel and Pletcher labs pioneered UV-Visible spectroscopy assays with simple food dye tracers to measure how much food was consumed based on the concentration of dye accumulated in adult flies (Shell et al., 2018, 2021; Shell & Grotewiel, 2022). Together, these assays in larvae could measure how feeding and locomotion are modified with different doses of antidepressants, and could be combined with direct electrochemical detection of serotonin to ascertain how neurochemistry relates to behavior in *Drosophila*.

Electrochemical techniques have been used to measure rapid serotonin changes with SSRIs and psychedelics in *Drosophila* and mice (Bowman et al., 2020; Dunham & Venton, 2020, 2022; Hashemi et al., 2009, 2012; Saylor et al., 2019; Wood et al., 2014). The Daws group used chronoamperometry to measure serotonin in SERT double knockout (-/-) mice and found that high doses of ketamine (32 mg/kg) slowed serotonin reuptake (Bowman et al., 2020), which led them to conclude that mSERT inhibition was required for ketamine’s antidepressant effects. However, they only measured serotonin changes with a single, high dose and did not investigate lower doses similar to a microdose treatment. Also in mice, the Hashemi lab used fast-scan cyclic voltammetry (FSCV) to show SSRIs, like escitalopram, citalopram, and fluoxetine, bind to SERT to inhibit serotonin reuptake (Hashemi et al., 2012; Ribaudo et al., 2021; Wood et al., 2014; Wood & Hashemi, 2013). Recently, our lab investigated optogenetically-stimulated serotonin release with several SSRIs using FSCV in *Drosophila* larvae and found concentration and reuptake changes that were similar to these in *vivo* responses in mice (Dunham & Venton, 2022; Ribaudo et al., 2021; Wood et al., 2014, p. 201; Wood & Hashemi, 2013). Additionally, SSRIs differentially affected serotonin, since some had more effects on increasing release while others only inhibited dSERT to slow reuptake (Dunham & Venton, 2022; Ribaudo et al., 2021; Stahl, 1998; Zhong et al., 2012). FSCV has also been used to measure serotonin changes with cocaine and phencyclidine (PCP) with *Drosophila* larvae (Borue et al., 2009, 2010; Pörzgen et al., 2001), however ketamine has not been explored. Thus, FSCV could be used to compare ketamine to SSRIs in *Drosophila* to clarify how serotonin changes at different doses to explain their effects on behavior.

The goal of this study was to compare changes in serotonin, locomotion, and feeding behaviors in *Drosophila* larvae after feeding ketamine or SSRI antidepressants. Here, we found that feeding low, 1 mM doses of ketamine for 24 hours did not affect serotonin concentration or reuptake, but 10 mM ketamine increased serotonin concentrations without inhibiting reuptake. However, at a high, 100 mM dose, which had anesthetic properties, ketamine inhibited dSERT and increased serotonin concentrations. SSRIs slowed serotonin reuptake at all doses, but affected serotonin concentrations differently based on their dSERT affinities. Feeding low doses (≤ 10 mM) of ketamine, escitalopram, and fluoxetine also increased feeding and locomotion behavior, while higher doses (100 mM) of ketamine and fluoxetine decreased these behaviors. Since ketamine did not primarily work through dSERT and has other targets, we also compared behavior changes with other NMDA receptor antagonists and 5-HT_1A and 2_ agonists and found NMDA receptor antagonism increases feeding, while serotonin receptor agonism increases locomotion. Thus, microdosing ketamine does not affect serotonin release or reuptake, although it does affect behavior, and may use different mechanisms than SSRIs. Ultimately, our work shows that *Drosophila* is a good model to determine different antidepressant mechanisms, and other genetic applications should be further explored to understand serotonin and behavior effects with ketamine and SSRIs.

## Methods

### Chemicals

Serotonin hydrochloride (CAS Number: 153-98-0), ketamine hydrochloride (1867-66-9), fluoxetine hydrochloride (56296-78-7), escitalopram oxalate (219861-08-2), paroxetine hydrochloride (110429-35-1), MK-801 hydrogen maleate (77086-22-7), 8-hydroxy-2-(di-n-propylamino) tetralin hydrobromide (8-OH-DPAT, 87394-87-4), buspirone hydrochloride (1078-80-2) and all-trans retinal (116-31-4) were purchased from Sigma Aldrich (St Louis, MO, USA).(Dunham & Venton, 2020, 2022) meta-Chlorophenylpiperazine (m-CPP, 65369-76-8) was purchased from Alfa Aesar (Haverhill, MA, USA). FD&C Blue No. 1 food dye powder (3844-45-9, Flavors and Color, Diamond Bar, CA, USA) was purchased on Amazon. For pre- and post-calibrations, a 1 mM stock solution of serotonin was prepared in 0.1 M HClO_4_. A final working solution of 1 µM serotonin was prepared by diluting the stock in phosphate buffer saline (PBS, 131.25 mM NaCl, 3.00 mM KCl, 10 mM NaH_2_PO_4_, 1.2 mM MgCl_2_, 2.0 mM Na_2_SO_4_, and 1.2 mM CaCl_2_ with the final pH adjusted to 7.4 with 1 M NaOH) (Dunham & Venton, 2020, 2022). Drugs were prepared in PBS and made fresh daily (Dunham & Venton, 2020, 2022).

### Microelectrode preparation

CFMEs were prepared as previously described (Dunham & Venton, 2020, 2022; Puthongkham et al., 2019). Briefly, a T-650 carbon fiber (Cytec Engineering Materials, West Patterson, NJ, USA) with a 7 µm diameter was aspirated into a standard 1.28 mm inner diameter x 0.69 mm outer diameter glass capillary tube (A-M Systems, Sequim, WA) with a vacuum pump. A capillary was then pulled by a Flaming Brown micropipette horizontal puller (Sutter Instrument, Novato, CA) to make two electrodes (Dunham & Venton, 2022, p. 202). Fibers were cut to 25-75 µm and epoxied by dipping the tip of the electrode into a solution of 14% m-phenylenediamine hardener (108-45-2, Acros Organics, Morris Plains, NH, USA) in Epon Resin 828 (25068-38-6, Miller Stephenson, Danbury, CT, USA) at 80–85 °C for 35 seconds. The CFMEs were cured at 100°C overnight and 150°C for at least 4 hours the next day.

### Electrochemical instrumentation

Electrochemical experiments were performed using a two-electrode system with a CFME working electrode backfilled with 1 M KCl (Dunham & Venton, 2020, 2022; Puthongkham et al., 2019). All potential measurements are reported versus a chloridized, Ag/AgCl wire reference electrode. Experiments were conducted in a covered, grounded Faraday cage to block out light. Before experiments, electrode tips were soaked in isopropyl alcohol for at least 10 minutes to clean the surface. The extended serotonin waveform (ESW, 0.2 V, 1.3 V, -0.1 V, 0.2 V, 1000 V/s) was continuously applied to electrodes using a WaveNeuro system (Pine Research, Durham, NC, USA) (Dunham & Venton, 2020, 2022). Data were collected with HDCV Analysis software (Department of Chemistry, University of North Carolina at Chapel Hill, USA). A flow-injection system with a six-port loop injector and air actuator (Valco Instruments, Houston, TX, USA) was used to pre- and post-calibrate CFMEs for in vitro experiments. PBS buffer was flowed at 2 mL/min using a syringe pump (Harvard Apparatus, Holliston, MA, USA) through a flow cell with the CFME tip inserted in solution. For calibration, 1 µM serotonin was injected for 5 seconds to determine current response. The concentration of serotonin released during *in vitro* experiments was determined using this calibration factor.

### Ventral nerve cord tissue preparation for optogenetic in vitro experiments

Methods for larva VNC dissection were previously described (Borue et al., 2009, 2010; Dunham & Venton, 2020, 2022; Privman & Venton, 2015). UAS-CsChrimson (Stockline BL#55136, Bloomington Drosophila Stock Center, Bloomington, IN, USA) virgin females were crossed with trh-Gal4 (BL#38389) flies and progeny heterozygous larvae were kept in the dark and raised on standard food (Food “J” for Janelia, Lab Express, Ann Arbor, MI, USA) mixed 250:1 with 100 mM all-trans retinal (ATR) until drug feeding was applied (Dunham & Venton, 2020, 2022). The ventral nerve cords (VNCs) of third instar “wandering” larvae were dissected in PBS kept on ice. A VNC was placed in an uncoated Petri dish dorsal side down containing 3 mL of room temperature PBS. A small slice of the lateral optic lobe was removed using the tip of a 22-gauge hypodermic needle. A CFME was implanted from the lateral edge of the tissue into the dorsal medial protocerebrum using a micromanipulator. Dissection and electrode insertion were performed under low light conditions to reserve the pools of releasable serotonin (Borue et al., 2010). CFMEs were allowed to equilibrate for 10 minutes in tissue in the dark prior to data collection (Dunham & Venton, 2020, 2022). Institutional ethical approval was not required for this study. Randomization procedures were not applied for allocation of different treatments. No exclusion criteria for samples were predetermined. No blinding was performed during data analysis.

### Optogenetic serotonin release

Optogenetic release of serotonin was stimulated by activating CsChrimson ion channels with red light from a 617 nM fiber-coupled high-power LED with a 200 µm core optical cable (ThorLabs, Newton, NJ, USA) (Borue et al., 2009, 2010; Dunham & Venton, 2020, 2022; Privman & Venton, 2015). A micromanipulator was used to center the fiber above the VNC tissue, and transistor-transistor logic (TTL) inputs to a T-cube LED controller (ThorLabs) were connected to the FSCV breakout box to control the light. TTL input was driven by electrical pulses from the WaveNeuro system and HDCV software to control frequency, pulse width, and number of pulses. For *in vitro* ketamine dose-response experiments, 120 biphasic pulses were delivered at 60 Hz with pulse width of 4 ms. An initial stimulation was recorded and 1 mL of the drug was slowly added to the Petri dish to not move the tissue or CFME (Dunham & Venton, 2022). Stimulations were repeated every 5 minutes for 30 minutes after a drug was added to allow the releasable pool of serotonin to replenish itself. For antidepressant feeding experiments for 24 hours, 30 biphasic pulses were applied with the same parameters.

### Feeding antidepressants to larvae for FSCV optogenetic experiments

Heterozygous progeny larvae (trh-Gal4; UAS-CsChrimson) were collected on day 5 after eating standard food mixed 250:1 with 100 mM ATR since birth. A stock drug solution that was 2x as concentrated as the final desired dose was made in PBS. Larvae were scooped out with food and gently mixed with an equal volume of drug. ATR was then added again in the same 250:1 ratio. Larvae were allowed to eat the food with drug for 24 hours before FSCV experiments (Figure 1, *n* = 6 larvae). The control larvae were mixed with only PBS.

**Figure 1.**
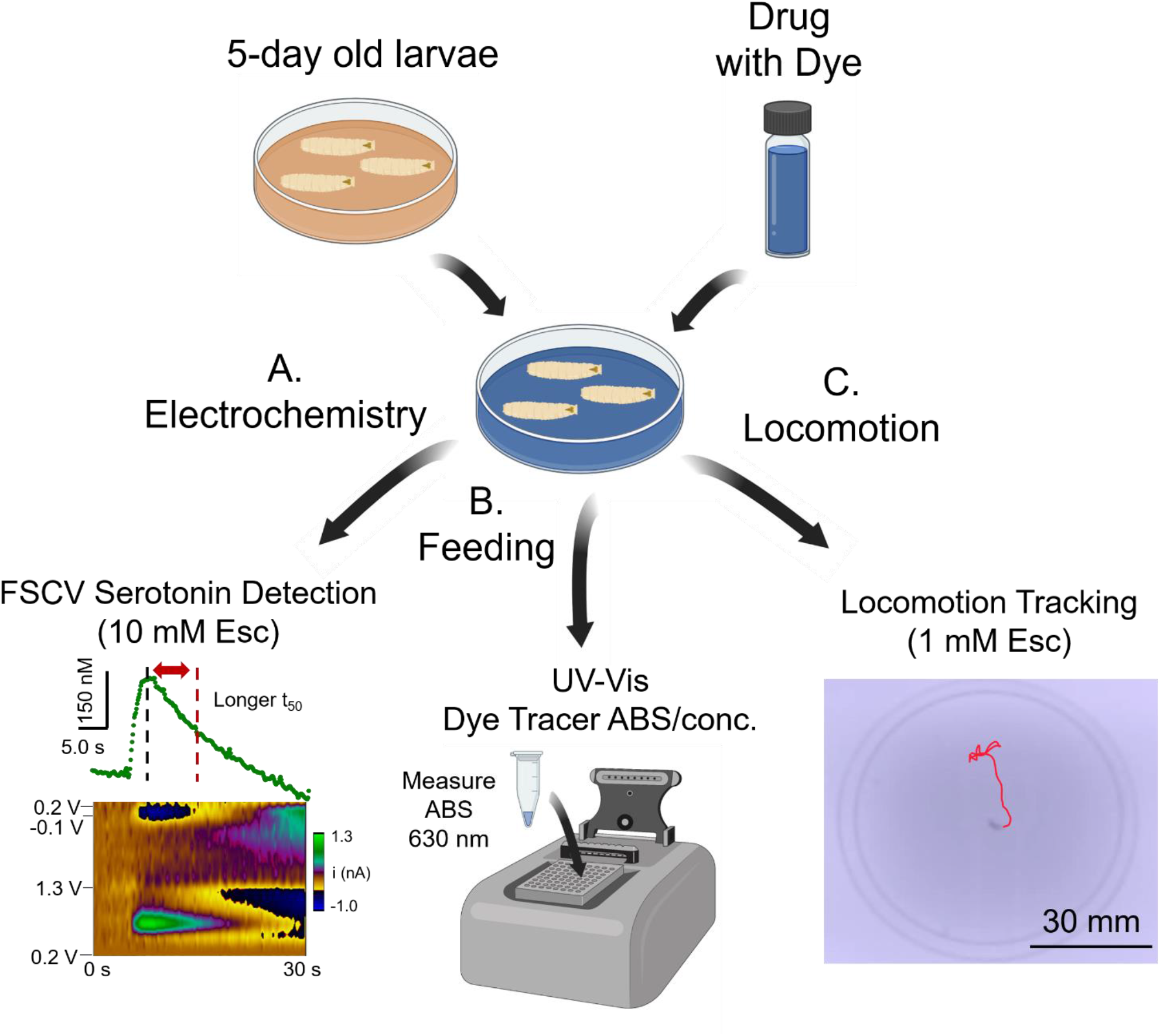
Feeding antidepressants to *Drosophila* larvae to measure real-time serotonin changes with feeding and locomotion behaviors. Larvae were collected on day 5 and ate antidepressants for 24 hrs. **A.** Fast-scan cyclic voltammetry (FSCV) and optogenetics were used to measure serotonin concentration and reuptake changes. Example FSCV conc. versus time and color plot show serotonin detected after feeding 10 mM escitalopram. Current was converted to serotonin conc. using a post-calibration factor, and reuptake is characterized by the time to half max decay (t_50_) of the peak detected in the trace (*n* = 6 larvae/ drug dose). **B.** Blue dye consumption was measured with a UV-Vis spectrophotometer from a tissue homogenate of *n* = 30 larvae. Blue dye absorbance was measured at 630 nm. **C.** Locomotion was tracked using PiVR and LoliTrack. A larva was placed on a 60 mm Petri dish. PiVR recorded a 60 sec video of the larva (*n* = 30 larvae/drug and dose), and LoliTrack was used to track their movement. Cartoons created in BioRender.

### UV-Vis dye tracer food consumption determination with different antidepressants

A 1% w/v FD&C Blue No. 1 solution was made by adding food dye powder to PBS. Methods are adapted from Shell et (Shell et al., 2018, 2021; Shell & Grotewiel, 2022). For drug feeding experiments, an antidepressant drug was added to the blue dye solution. On day 5, larvae were scooped with their food and mixed with the blue dye drug solution. Larvae ate the drug and blue dye tracer for 24 hours. After, 30 larvae were collected and placed into 500 µL of deionized (DI) water in an Eppendorf tube and centrifuged at 13,000 rpm for 30 mins using an AccuSpin Micro 17 Centrifuge (Fisher Scientific, Waltham, MA, USA). A micro spatula was used to crush the larvae into a tissue homogenate and it centrifuged for 15 mins. Tissue homogenates were discarded and 50 µL of the blue dye supernatant was added to 200 µL DI water to make a 1:5 dilution. The absorbance of the blue dye sample was measured at 630 nm using a Tecan plate reader Ultraviolet-Visible spectrophotometer (Tecan, Männedorf, Switzerland). For each drug, *n* = 5 samples (n = 30 larvae/sample) were collected and measured with 3 technical replicates. A calibration curve was constructed from 5 – 250 µM dye, and the slope of the calibration graph was used to determine the concentration of dye in the supernatant to calculate the mass of food eaten per larva per day.

### Real-time Drosophila larvae locomotion tracking

The Raspberry Pi based Virtual Reality system (PiVR, v. 3) was constructed following the methods in Tadres and Louis 2020 and the “Build your own PiVR” section on their website.(Tadres & Louis, 2020) The computer casing parts and camera tower were printed using a Stratasys F170 3D Printer (Strartasys, Eden Prairie, MN, USA) at UVA’s MAE Rapid Prototyping and Machine labs using the files from the Louis Lab GitLab. The camera resolution was set to 640×480, and the animal detection method was set for third instar *Drosophila* larvae. After eating a drug for 24 hours, a larva was placed in the center of a 60 mm Petri dish and their movements were recorded for 60 s (*n* = 30 larvae). The individual recording files were then converted from h2p6 format to MPEG4 format using File Viewer Plus (Sharpened Productions, Minneapolis, MN, USA, v. 4.3.). LoliTrack software (Loligo Systems, Viborg, Denmark, v. 5.2.0) was used to track the linear distance the larvae traveled from the video recordings using their “Tracking 2D” setting.

### Statistics and data analysis

Data are the mean ± the standard error of the mean (SEM) for *n* number of *Drosophila* larvae. For SSRI and ketamine feeding electrochemistry experiments, *n* = 6 larvae (larvae were not sexed, so both males and females were used) (Dunham & Venton, 2022). For drug dose response curve experiments, each dose is at least *n* = 4 larvae. For a sample calculation, we performed power analysis in MedCalc (MedCalc Statistical Software, Ostend, Belgium) using the paired sample t -test where ɒ = 0.05, 1-β = 0.1, mean difference in current detected = 0.45 nA, and standard deviation of differences = 0.2 nA, which results in *n* = 4 larvae (Dunham & Venton, 2022). Statistics were performed in GraphPad Prism 8.0 (GraphPad Software, La Jolla, CA). Data were normally-distributed for multiple drug comparisons (all KS distance ≥ 0.1980, Kolmogrorov-Smirnov, p ˃ 0.1000). Significance was determined at a 95% confidence level for One-Way ANOVA, Tukey’s post-hoc test, and comparison of fits. No test for outliers was performed.

## Results

### Ketamine does not affect serotonin at low doses, but blocks reuptake at higher doses after 24 hours

To characterize chronic consumption of antidepressants in *Drosophila*, we fed larvae several doses of ketamine and SSRIs for 24 hours and measured optogenetically-released serotonin with FSCV. Figure 2A-K shows serotonin concentration and reuptake changes with FSCV serotonin conc. versus *t* plots after feeding larvae (A) PBS only, (B) 1 mM ketamine, (C) 10 mM ketamine, (D) 100 mM ketamine, (E) 1 mM paroxetine, (F) 10 mM paroxetine, (G) 1 mM fluoxetine, (H) 10 mM fluoxetine, (I) 100 mM fluoxetine, (J) 1 mM escitalopram, and (K) 10 mM escitalopram. Ketamine did not affect serotonin concentration or reuptake at 1 mM, but serotonin concentrations did increase at 10 mM. However, ketamine both increased and blocked serotonin reuptake by inhibiting dSERT at 100 mM, which is illustrated by the longer t_50_ in Fig. 2D. At the low 1 mM dose, escitalopram and fluoxetine also did not increase serotonin concentrations, but serotonin reuptake was slower with both of these SSRIs. In comparison, paroxetine increased serotonin concentrations and reuptake at this low dose because of its high dSERT affinity.(Dunham & Venton, 2022) Because fluoxetine did not change serotonin release at lower doses, we fed a higher (100 mM) doses, which increased serotonin concentrations and slowed reuptake.

**Figure 2.**
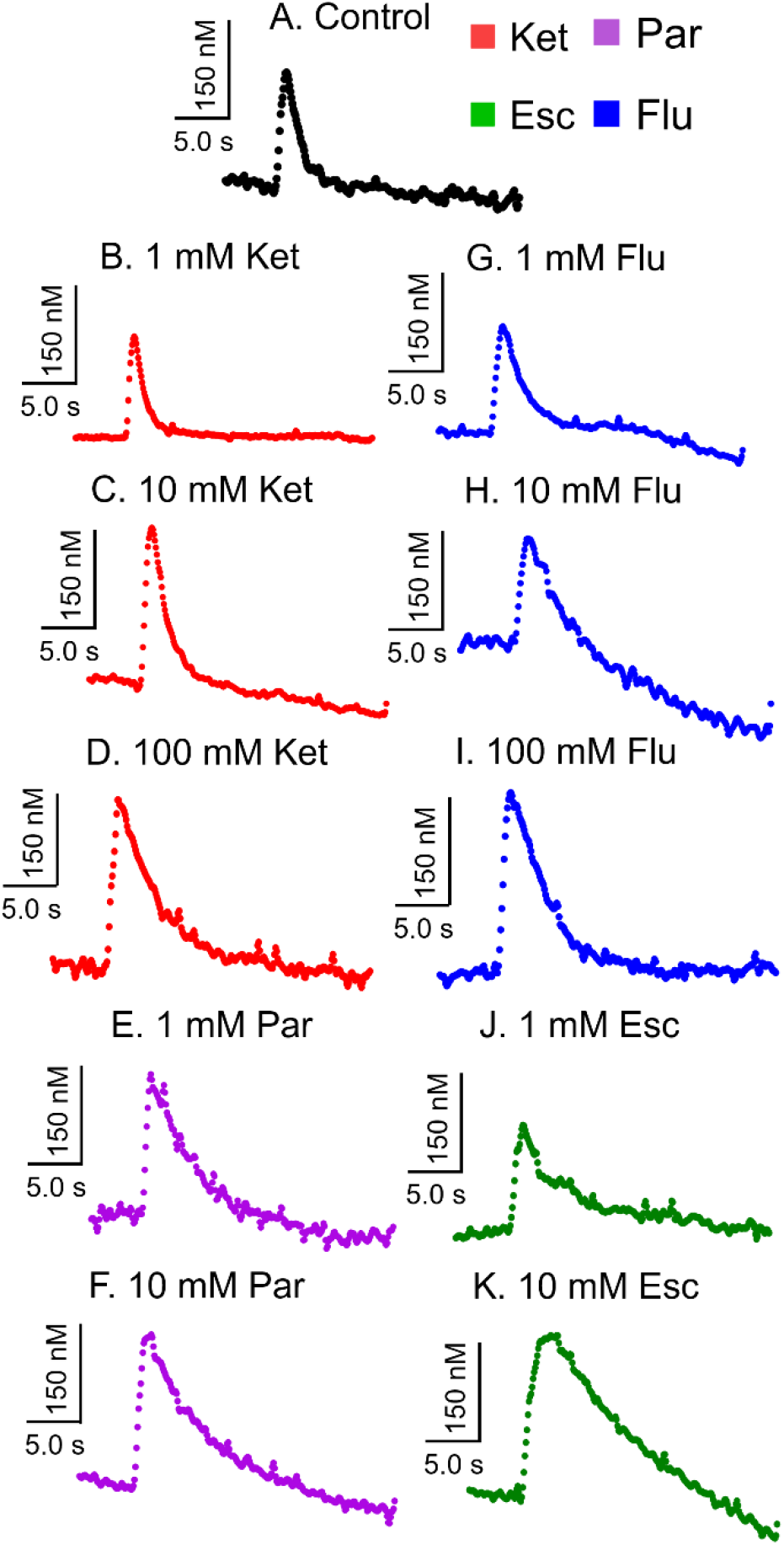
Feeding low (1 mM) and mid-doses (10 mM) ketamine does not inhibit serotonin (5-HT) reuptake, but does at high (100 mM) doses compared to SSRIs. *Drosophila* larvae were fed (**A**, black) PBS only, (**B**, red) 1 mM ketamine, (**C**) 10 mM ketamine, (**D**) 100 mM ketamine, (**E**, purple) 1 mM paroxetine, (**F**) 10 mM paroxetine, (**G**, blue) 1 mM fluoxetine, (**H**) 10 mM fluoxetine, (**I**) 100 mM fluoxetine, (**J**, green) 1 mM fluoxetine, and (**K**) 10 mM escitalopram for 24 hours (*n* = 6 larvae). Ketamine did not affect serotonin concentration or reuptake at 1 mM, but increased concentrations at 10 mM and slowed reuptake by inhibiting dSERT at 100 mM. At low doses, paroxetine only increased serotonin concentrations, but reuptake increased with paroxetine, escitalopram, and fluoxetine at low doses. At mid-doses, paroxetine and escitalopram increased concentration, but not fluoxetine. However, fluoxetine increased both serotonin concentration and reuptake at 100 mM.

Figure 3A-B shows serotonin concentration and reuptake changes with each antidepressant dose after feeding for 24 hours. One-Way ANOVA and Tukey’s post-hoc test showed that feeding antidepressants significantly affected serotonin concentration (Fig. 3A, F_(10,55)_ = 15.67, *p* ≤ 0.0001, *n* = 6) and reuptake (Fig. 3B, F_(10,55)_ = 188.1, *p* ≤ 0.0001, *n* = 6). With ketamine, serotonin concentrations did not increase with 1 mM, but increased with 10 and 100 mM (*p* ˂ 0.0001). Additionally, ketamine did not affect reuptake at 1 or 10 mM, but inhibited reuptake by blocking dSERT at 100 mM (*p* ˂ 0.0001). With SSRIs, reuptake was inhibited with every drug and dose (all *p* ˂ 0.0001). However, serotonin concentrations increased based on each SSRI’s individual dSERT affinity. For example, paroxetine increased serotonin concentrations at 1 mM (*p* ˂ 0.01) because of its high dSERT affinity, while escitalopram and fluoxetine did affect serotonin concentration. In addition to feeding ketamine, we also bath-applied ketamine directly to VNC tissue, which is illustrated in Figure S1. Here, there were similar trends as low doses (≤ 5 µM) did not affect serotonin release or reuptake, while higher doses (1 mM) blocked serotonin reuptake to increase serotonin concentrations. Together, these data show that ketamine, unlike SSRIs, does not affect serotonin release or reuptake at microdoses.

**Figure 3.**
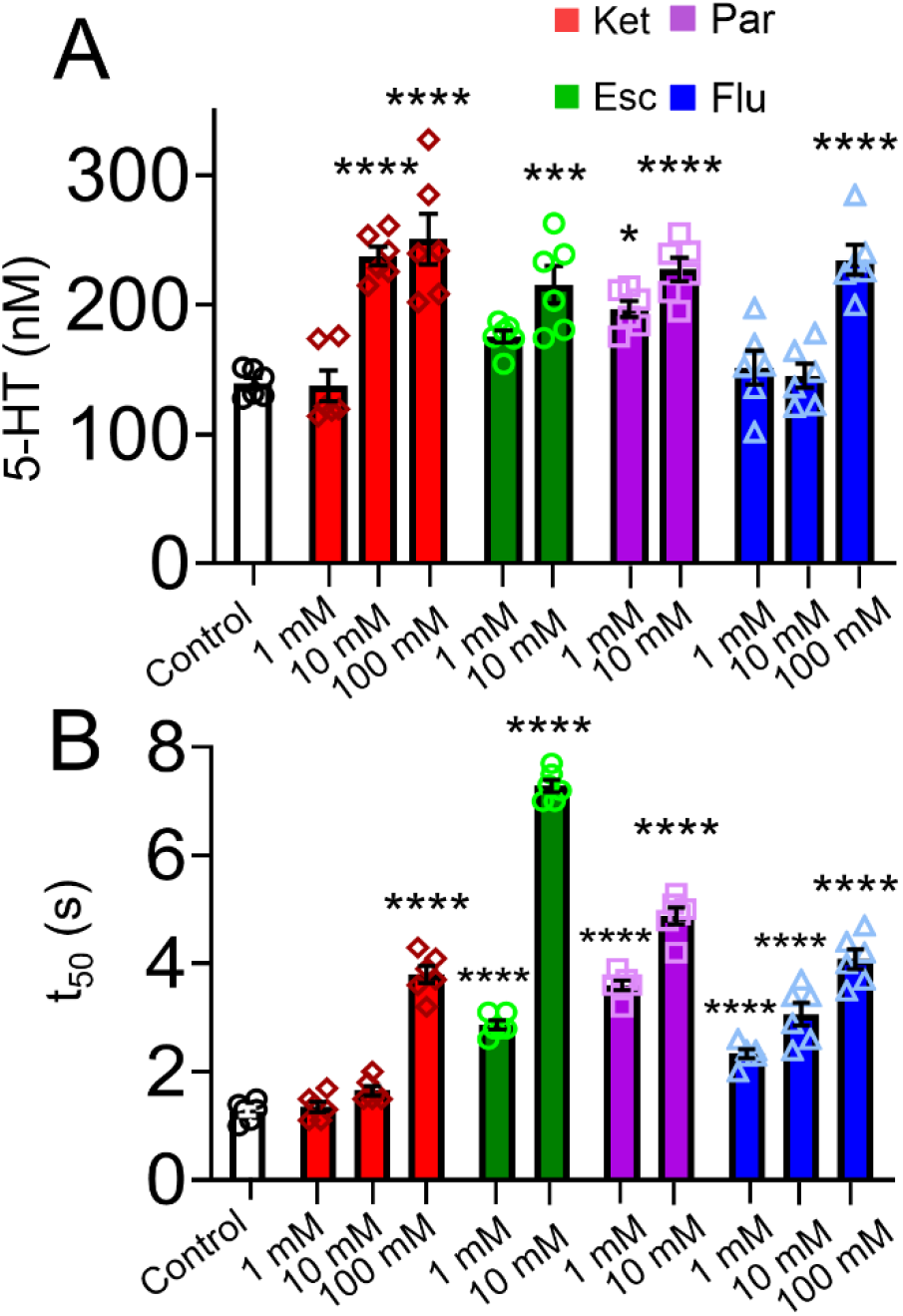
Ketamine does not affect serotonin at low, 1 mM doses, but increases serotonin concentrations and inhibits reuptake with a high, 100 mM dose. Meanwhile, SSRIs block reuptake at all doses. Changes in serotonin (**A**) concentration and (**B**) reuptake were measured using FSCV. One-Way ANOVA and Tukey’s post-hoc test showed significant effects of drug on serotonin concentration (F_(10,55)_ = 15.67, *p* ≤ 0.0001, *n* = 6) and reuptake (F_(10,55)_ = 188.1, *p* ≤ 0.0001, *n* = 6). At 1 mM, ketamine does not affect serotonin concentration or reuptake, but at 10 mM it increases serotonin concentrations (*p* ≤ 0.0001). However, ketamine inhibits serotonin reuptake (*p* ˂ 0.0001) and increases concentrations at 100 mM (*p* ≤ 0.0001). SSRIs increase serotonin reuptake at all doses (all *p* ˂ 0.0001), but show differences in serotonin concentration increases from their individual dSERT affinities.

### Low doses of ketamine and escitalopram increase feeding behaviors

A common symptom of depression and side effect of antidepressants is weight gain or loss (De Vry & Schreiber, 2000; Keszthelyi et al., 2009). To understand how SSRIs and ketamine affect feeding behaviors, we used blue food dye tracers with UV-Vis spectroscopy to measure the amount of food *Drosophila* larvae ate after 24 hours. A linear calibration curve of absorbance vs. dye concentration was made to calculate the dye consumed by a larva (Figure S2). Figure 4 compares the mass of food eaten (µg) per larva per day (*n* = 5 collection samples/dose with *n* =30 larvae/sample). Overall, there was a significant effect of antidepressant type and dose on food consumption (Fig. 4, One-Way ANOVA, F_(10,44)_ = 238.5, *p* ≤ 0.0001, *n* = 5), and Tukey’s post-hoc test showed that larvae significantly consumed more food with 1 mM ketamine, 1 mM escitalopram, and 10 mM ketamine (*\*\*\*\* p* ˂ 0.0001), but less food was consumed with 100 mM ketamine (*\* p* ˂ 0.05).

**Figure 4.**
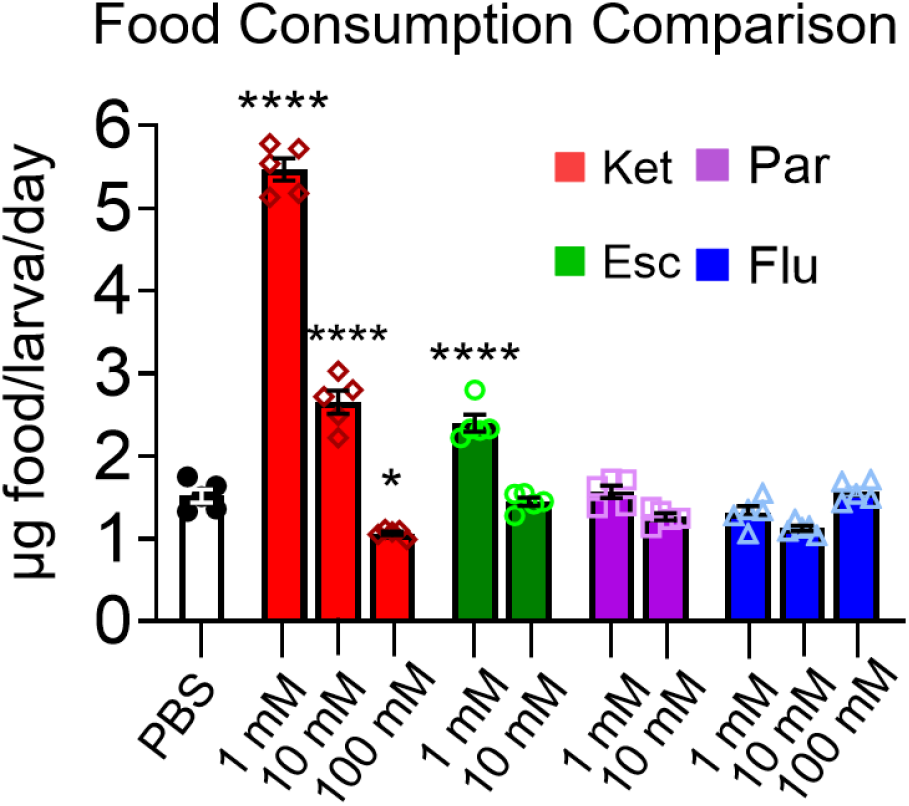
Food consumption for different antidepressant drugs (*n* = 5 samples/dose with *n* = 30 larvae/sample). One-Way ANOVA (F_(10,44)_ = 238.5, *p* ≤ 0.0001, *n* = 5) and Tukey’s post-hoc comparisons found the amount of blue food eaten significantly increased with 1 mM ketamine, 10 mM ketamine, and 1 mM escitalopram (all \*\*\*\**p*< 0.0001). However, the amount of food consumed significantly decreased with 100 mM ketamine (*p* = 0.0134).

### Low doses of ketamine, escitalopram, and fluoxetine increase locomotion behaviors

In addition to changes in appetite and feeding, antidepressants have been previously shown to affect locomotion (Hidalgo et al., 2017; Majeed et al., 2016; Silva et al., 2014). Here, we used PiVR to record *Drosophila* larvae and LoliTrack tracking software to characterize locomotion changes after feeding ketamine and SSRIs for 24 hours. Here, we measured the linear distance traveled from the center of the petri dish, and disregarded sweeping head and flopping larvae movements. Figure 5A-E shows still images with the path the larva traveled drawn in red after eating food mixed with (A) PBS only, (B) 1 mM ketamine, (C) 1 mM escitalopram, (D) 1 mM paroxetine, and (E) 1 mM fluoxetine. Additionally, Figure S3 shows still images of larvae at the other doses tested. One-Way ANOVA showed there was a significant overall effect of antidepressant on distance traveled (Fig. 5F, F_(10,319)_ = 72.35, *p* ≤ 0.0001, *n* = 30) and Tukey’s post-hoc test showed that distance traveled significantly increased with 1 mM ketamine, 1 mM escitalopram, 1 mM fluoxetine, and 10 mM fluoxetine (all *p\*\*\*\** ˂ 0.0001). However, distance traveled significantly decreased with 100 mM fluoxetine (*p\*\** ˂ 0.01).

**Figure 5.**
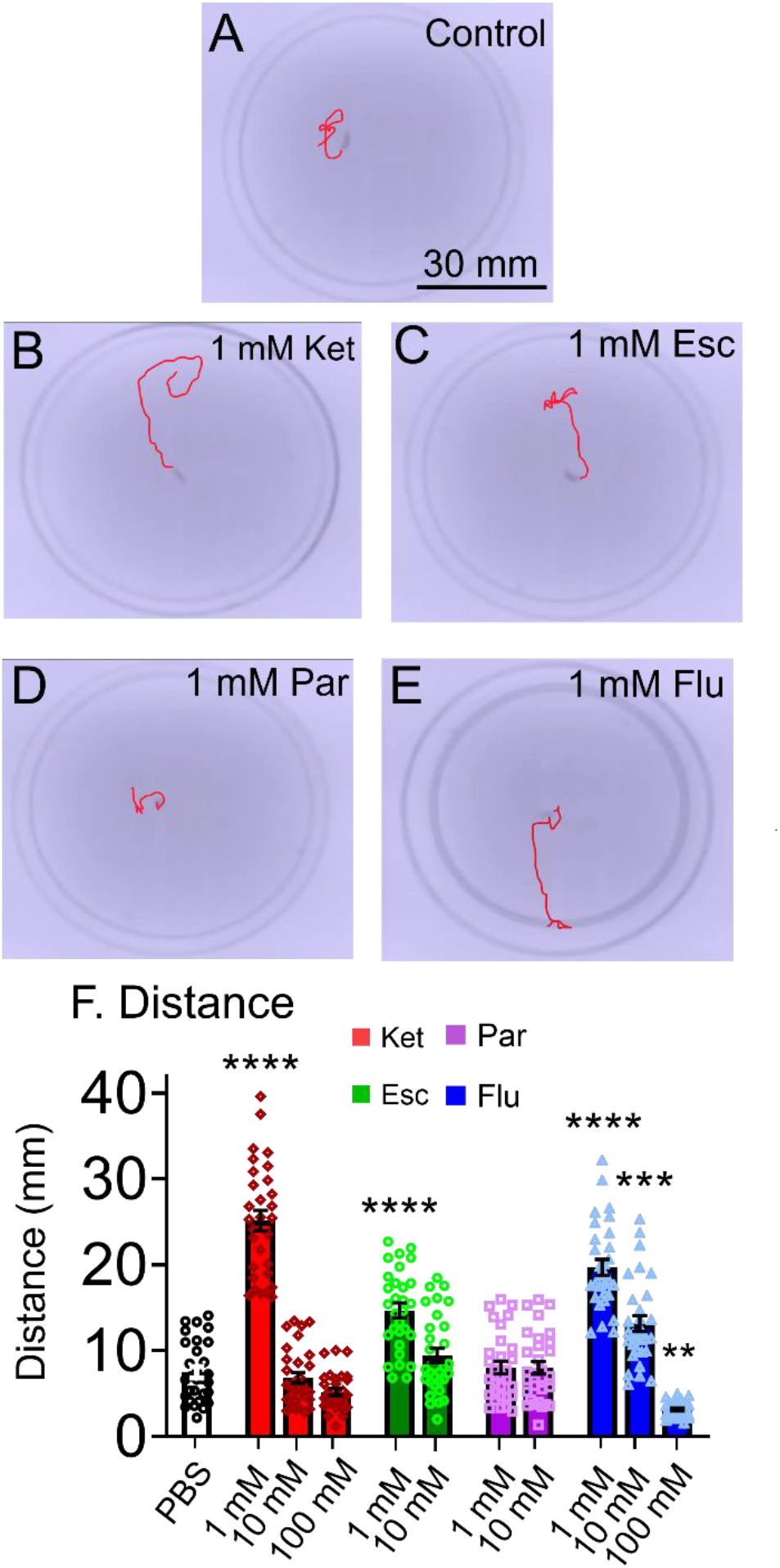
Locomotion tracking comparison after *Drosophila* larvae ate different antidepressant drugs for 24 hours (*n* = 30 larvae/dose). A larva’s locomotion was tracked for 60 s after feeding (A) PBS, (B) 1 mM ketamine, (C) 1 mM escitalopram, (D) 1 mM paroxetine, and (E) 1 mM fluoxetine. Larvae showed increased locomotion with 1 mM ketamine, 1 mM escitalopram, 1 mM fluoxetine, and 10 mM fluoxetine. However, locomotion decreased with 100 mM fluoxetine. F. One-Way ANOVA (F_(10,319)_ = 72.35, *p* ≤ 0.0001, *n* = 30) and Tukey’s post-hoc comparisons found the distance traveled significantly increased with 1 mM ketamine, 1 mM escitalopram, 1 mM fluoxetine, and 10 mM fluoxetine (all p****< 0.0001). However, distance significantly decreased with 100 mM fluoxetine (p ≥ 0.0020).

### NMDA receptor antagonist increase feeding at low doses, while 5-HT receptor agonists increase locomotion

Our data indicate that ketamine does not primarily work through SERT and may act via a different mechanism. Ketamine is classified as an NMDA receptor antagonist and also activates serotonin receptors 5-HT_1A, 2A, and 2C_ Kraus et al., 2019; Moghaddam et al., 1997). To understand how these specific receptors affect locomotion and feeding compared to ketamine and SSRIs, we fed a low dose (1 mM) MK-801, an NMDA receptor antagonist (Verma & Moghaddam, n.d.), 8-OH-DPAT, a full 5-HT_1A_ agonist, and buspirone, a partial 5-HT_1A_ agonist (De Boer & Koolhaas, 2005; Gould et al., 2011; Johnson et al., 2009), and m-CPP, a psychoactive stimulant that acts as a 5-HT_2_ agonist in *Drosophila* (De Vry & Schreiber, 2000; Majeed et al., 2016). Figure 6A compares the food eaten per larva per day with low doses of all drugs (*n* = 5 collection samples/dose with *n* = 30 larvae/sample). Overall, there was a significant effect of drug on food consumption (Fig. 6A, One-Way ANOVA, F_(8,36)_ = 252.8, *p* ≤ 0.0001, *n* = 5), and Tukey’s post-hoc test showed that low dose MK-801 (*p* ˂ 0.0001) and 8-OH-DPAT (*p* ˂ 0.001) significantly increased feeding, similar to ketamine. However, feeding also increased with 8-OH-DPAT for 5-HT_1A_, which is similar to the increases in feeding with escitalopram. Figure 6B also shows the distance traveled after eating 1 mM of the different drugs. Images of larvae locomotion paths can be found in Figure S4. One-Way ANOVA showed there was a significant overall effect of drug on distance traveled (Fig. 8B, F_(8,261)_ = 80.72, *p* ≤ 0.0001, *n* = 30) and Tukey’s post-hoc test showed that distance traveled dramatically increased with 1 mM 8-OH-DPAT, m-CPP, and buspirone (all *p* ≤ 0.0001), which are similar to the changes for 1 mM escitalopram and fluoxetine. However, distance only slightly increased with MK-801 (*p* ˂ 0.1). Together, these data suggest that NMDA receptor antagonism increases feeding, while serotonin receptor agonism increases locomotion, which could explain these behaviors in ketamine and SSRIs.

**Figure 6.**
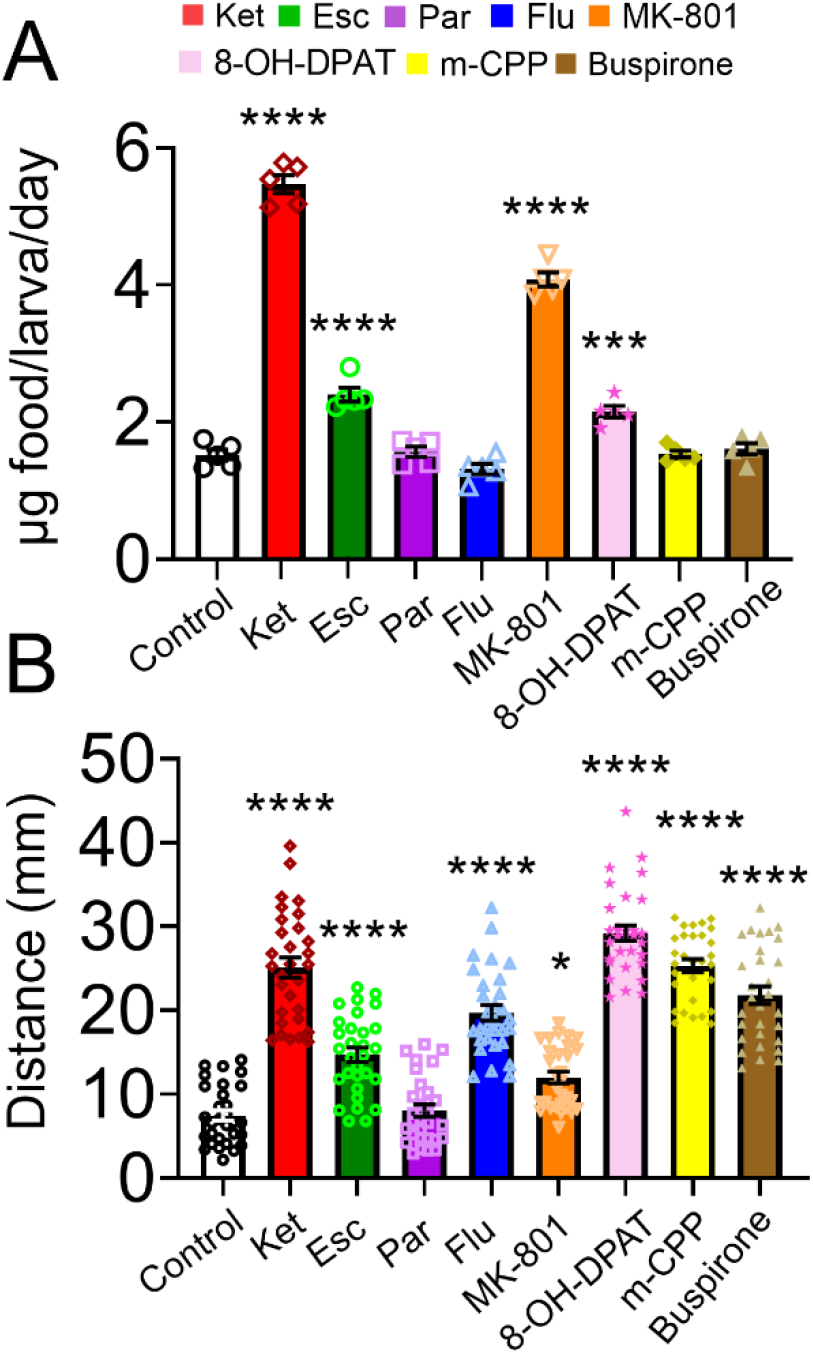
Feeding and locomotion changes after *Drosophila* larvae ate low doses (1 mM) of receptor specific drugs drugs for 24 hours compared to ketamine and SSRIs (*n* = 30 larvae/drug). MK-801 is a NMDA antagonist, 8-OH-DPAT a 5-HT_1A_ agonist, m-CPP a 5-HT_2B_ partial agonist, and buspirone a 5-HT_2C_ agonist. A. Food consumption comparison. One-Way ANOVA (F_(8,36)_ = 252.8, *p* ≤ 0.0001, *n* = 5) and Tukey’s post-hoc comparisons found the amount of blue food eaten significantly increased with 1 mM ketamine, escitalopram, and MK-801 (all \*\*\*\**p*< 0.0001), as well as with 1 mM 8-OH-DPAT (\*\*\**p*< 0.001). B. Locomotion was tracked for 60 s with the other receptor drugs versus ketamine and SSRIs. One-Way ANOVA (F_(8,261)_ = 80.72, *p* ≤ 0.0001, *n* = 30) and Tukey’s post-hoc comparisons found the distance traveled significantly increased with 1 mM ketamine, escitalopram, fluoxetine, 8-OH-DPAT, m-CPP, and buspirone (all ****p< 0.0001), but only slightly increased with 1 mM MK-801 (p*< 0.1).

## Discussion

Our results show that feeding ketamine to *Drosophila* larvae does not affect serotonin at low microdoses, but increases serotonin release at mid-doses, and inhibits serotonin reuptake at higher, anesthetic doses. In contrast, all SSRI doses slowed serotonin reuptake, but they affected serotonin concentrations differently based on their dSERT affinities. Interestingly, we saw similar dose-dependent effects on behavior where low doses of ketamine, escitalopram, and fluoxetine increased feeding and locomotion behaviors, while higher doses of ketamine and fluoxetine decreased them. Together, these electrochemical and behavioral data indicate that dSERT inhibition is not required for ketamine to increase feeding and locomotion. Additional pharmacology behavior experiments also showed that NMDA receptor antagonism increases feeding, while 5-HT_1A_ and 5-HT_2_ agonism increases locomotion, which may be targets that ketamine affects in order to change behavior. Altogether, future studies could investigate the extent to which ketamine works through these receptors in *Drosophila*, but this work shows that *Drosophila* is a good model to rapidly study pharmacological mechanisms of ketamine and other antidepressants, because neurochemical changes can be compared to behavioral assays.

### Ketamine does not affect SERT at low doses, in contrast to SSRIs

Here, we fed larvae different doses of SSRIs and ketamine for 24 hours to compare long-term feeding effects. For ketamine, the lowest, 1 mM dose did not affect serotonin, while a 10 mM mid-dose only increased serotonin concentrations and did not inhibit reuptake. However, after feeding larvae 100 mM ketamine, serotonin concentrations dramatically increased and reuptake slowed by inhibiting dSERT. We also saw similar serotonin release and reuptake changes when we bath-applied ketamine directly to the larva VNC tissue (Fig. S1), where there were no effects on serotonin at low doses ≤ 5 µM, and concentration and reuptake significantly increased at the highest 1 mM dose. Although we cannot determine how much ketamine or SSRI is taken up into the VNC tissue, the feeding doses are similar to doses that were 100-fold lower when bath applied, which means the VNCs likely adsorb micromolar concentrations of drug (Dunham & Venton, 2022). Our data also shows that ketamine does not change serotonin concentrations or reuptake at low doses, which would be similar to the therapeutic doses used for microdosing for antidepressant effects. In contrast to ketamine, all SSRIs affected uptake at low doses and varied serotonin concentrations based on their dSERT affinities. For example, fluoxetine and escitalopram slowed serotonin reuptake at the low feeding doses, but only increased release at higher doses (Dunham & Venton, 2022), while 1 mM paroxetine showed high concentration changes and slower reuptake, which was similar to the higher doses we previously bath-applied (Dunham & Venton, 2022). Ultimately, these data suggest ketamine does not have similar effects on SERT compared to SSRIs at low doses.

The effects of ketamine on the serotonin system have been studied in small mammals and humans, but how microdoses affect this system is ambiguous (West et al., 2023). For instance, some literature suggests microdosing ketamine relies on SERT inhibition to increase serotonin concentrations after 24 hours (Kraus et al., 2019; Li, 2020; Pham et al., 2017; Spies et al., 2018), while other studies suggest that immediate responses (≤ 24 hours) may rely more on serotonin (5-HT_1A,2A,2C_) receptor activation to increase serotonin concentrations. However, another study with PET imaging in humans saw that SERT was 70-80% occupied by various SSRIs, but not affected by low doses of ketamine (Spies et al., 2018). Here, the authors also suggested SERT may be inhibited by higher, anesthetic ketamine doses, which is in agreement with our data in *Drosophila*. Furthermore, Can et al. used binding affinity assays to determine ketamine did not bind to SERT, norepinephrine, or dopamine transporters at concentrations ≤ 10 µM, which is similar to our bath application data for SERT (Can et al., 2016). In mice, ketamine also does not change dopamine at microdoses, but did cause hyperlocomotion and increased concentrations at higher anesthetic doses (Can et al., 2016). Additionally, the Daws group compared serotonin changes in real-time with wild type (WT) mice versus SERT double knockout (-/-) mice after injecting a high dose of 32 mg/kg ketamine (Bowman et al., 2020). This high ketamine dose blocked reuptake but had no effect on serotonin concentrations in the WT mice, and the SERT -/-mice were not affected. The Daws group also tried a lower, 3.2 mg/kg ketamine dose and found no changes in locomotion, which is similar to our 10 mM feeding dose data. Overall, our results in *Drosophila* were similar to this mammalian data as serotonin reuptake only significantly increased with higher doses. Thus, these results provide a more comprehensive dose response in *Drosophila*, which shows that feeding low doses for 24 hours does not affect SERT, but feeding higher, anesthetic doses blocks SERT. Ultimately, SERT blockade does not seem responsible for behavioral changes caused by feeding microdoses of ketamine.

### Ketamine and SSRIs change feeding and locomotion similarly, even though serotonin changes are different

Common symptoms of depression include weight gain or loss from changes in appetite and activity, which can be affected by taking antidepressants (Knapp et al., 2022; Majeed et al., 2016). However, it is not understood whether these behaviors are caused by serotonin changes. Here, we measured feeding and locomotion changes after feeding wildtype larvae ketamine and SSRIs for 24 hours. Primarily, we saw the low doses (1 mM) of ketamine, escitalopram, and fluoxetine increased both feeding and locomotion (Figure 4 and 5). Although the effects of ketamine on feeding had not been characterized in *Drosophila*, several new clinical studies have shown that microdosing ketamine increases appetite and has positive effects for patients with co-morbid eating disorders and depression (Ragnhildstveit et al., 2022; Robison et al., 2022; Schwartz et al., 2021). Similar to our *Drosophila* studies, low doses of ketamine also cause hyperlocomotion in mammals (Can et al., 2016; Irifune et al., 1991). With SSRIs, Hidalgo *et al*. investigated locomotion changes after feeding *Drosophila* larvae a high dose of fluoxetine (Hidalgo et al., 2017), and found that fluoxetine decreased locomotion, which is similar to our 100 mM fluoxetine data. Thus, *Drosophila* behavioral data matches with mammalian data and it is a good model organism to study these effects.

These results show that antidepressants and ketamine change feeding and locomotion behaviors in a dose-dependent manner. However, while behavioral changes are similar between SSRIs and ketamine, the serotonin changes were not similar. For instance, 1 and 10 mM ketamine, as well as 1 mM escitalopram increased feeding. At 1 mM, escitalopram inhibits dSERT, but ketamine does not. Interestingly, only a high ketamine dose inhibited dSERT and that dose decreased feeding, likely due to its anesthetic effects. Thus, the feeding effects of ketamine are not due to SERT inhibition. Likewise, we saw similar effects with locomotion, where low, 1 mM doses of ketamine, escitalopram, and fluoxetine increased locomotion, but serotonin concentrations were not significantly different. However, escitalopram and fluoxetine both significantly slowed reuptake, unlike ketamine. Similar to the feeding data, locomotion decreased with higher doses of ketamine and fluoxetine, even though serotonin concentrations increased and reuptake was inhibited. The Daws group found with their mammalian data that only the highest ketamine dose (32 mg/kg) decreased locomotion, which is similar to our data (Bowman et al., 2020). Thus, there is no correlation between increased serotonin or SERT inhibition and increased feeding and locomotion after ketamine, which suggests SERT inhibition is not the primary mechanism controlling these antidepressant behaviors. It is important to also note that only low doses of ketamine produce antidepressant effects because higher doses act as an anesthetic (Kraus et al., 2019; Stahl, 2013; Zanos et al., 2018).

### The antidepressant mechanisms of ketamine may rely on NMDA and serotonin receptors

Ketamine has effects on several neuromodulatory systems (Kraus et al., 2019). Formally, ketamine is described as an NDMA receptor antagonist, which inhibits the glutamatergic system (Kraus et al., 2019; Moghaddam et al., 1997; Verma & Moghaddam, n.d.). Although glutamate is an excitatory molecule in mammals (Galvanho et al., 2020), it shows excitatory and inhibitory effects in *Drosophila* in their neuromuscular junction (Zimmerman et al., 2017) and olfactory system (Liu & Wilson, 2013), respectively. It also acts as a 5-HT_1A, 2A, and 2C_ receptor agonist, which activates serotonin neurons,(Kraus et al., 2019) and *Drosophila* show homology for these receptors (Majeed et al., 2016; Silva et al., 2014). Since ketamine has several effects on different neurotransmitter systems, we further tested the behavioral effects of other drugs with high NMDA, 5-HT_1A_, and 5-HT_2_ receptor affinity. Our main findings were that the NMDA antagonist, MK-801, and 5-HT_1A_ agonist, 8-OH-DPAT, increased feeding, while 8-OH-DPAT, buspirone (5-HT_1A_ partial agonist), and m-CPP (5-HT_2_ agonist) increased locomotion. Mammalian studies are clear that SERT is required for antidepressants effects of SSRIs (Hashemi et al., 2012; Ribaudo et al., 2021; Saylor et al., 2019; Wood et al., 2014; Wood & Hashemi, 2013), but different behavioral effects with microdoses may be due to other targets as well. In rats, escitalopram activates 5-HT_1A_ receptors (El Mansari et al., 2005; Fakhoury, 2016), and our data in flies show that low doses of escitalopram increase both locomotion and feeding behavior, similar to our results with the 5HT_1A_ agonist, 8-OH-DPAT. Additionally, fluoxetine blocks 5-HT_2C_ receptors at higher doses (Ni & Miledi, 1997), which could cause the decreased locomotion we observed at these doses, since a 5HT_2_ agonist (m-CPP) increased locomotion. Altogether, more studies could examine these other targets for SSRIs and ketamine to determine the mechanisms of these behavioral effects.

This study shows that SERT inhibition is not required to achieve behavioral effects with microdoses of ketamine, and that ketamine may increase feeding behavior via NMDA receptor antagonism, while locomotion behavior may be caused by ketamine acting as a 5-HT_1A_ or 5-HT_2_ agonist (Kraus et al., 2019; Spies et al., 2018). Future studies in genetically altered flies, where NMDA or serotonin receptors are knocked-down, would help confirm these are the targets of ketamine or other SSRIs to elicit behavioral effects (Shin et al., 2020; Silva et al., 2014). Future work should also investigate the real-time changes of microdoses of ketamine or SSRIs on other neurotransmitters, such as glutamate (Marvin et al., 2018; Richter et al., 2018). However, this work shows that *Drosophila* is a good model to rapidly characterize neurotransmitter and behavior changes with different antidepressants, such as ketamine versus SSRIs, and is useful for discerning mechanisms of neurotransmitter control of behavior.

### Conclusions

Overall, this work shows that feeding microdoses of ketamine to Drosophila larvae does not inhibit SERT, unlike SSRIs, but does cause increased locomotion and feeding. At higher doses (100 mM), however, ketamine inhibited dSERT to increase serotonin and decreased locomotion and feeding from its anesthetic effects. Low doses (≤ 10 mM) of escitalopram and fluoxetine also increased feeding and locomotion behaviors, but mainly increased reuptake without increasing serotonin. Feeding a NMDA antagonist evoked similar increases in feeding to ketamine, while feeding 5HT_1A_ and 5HT_2_ agonists increased locomotion, similar to ketamine. Altogether, these data demonstrate that microdosing ketamine works through a different mechanism than SSRIs for behavioral effects. Ultimately, *Drosophila* is a good model to discern antidepressant and behavioral mechanisms with ketamine and SSRIs and future studies should investigate how they change with other neurotransmitters, such as glutamate, using genetic knockdowns of NMDA and serotonin receptors.

## Abbreviations

(ATR): all-trans retinal
(CFME): carbon fiber microelectrode
(DI): deionized water
(dSERT): *Drosophila* serotonin transporter
(Esc): escitalopram
(ESW): extended serotonin waveform
(FSCV): fast-scan cyclic voltammetry
(Flu): fluoxetine
(FDA): Food & Drug Administration
(iGluSnFR): Glutamate-sensitive fluorescent reporters
(Ket): ketamine
(MDD): major depressive disorder
(m-CPP): meta-Chlorophenylpiperazine
(Par): paroxetine
(PCP): phencyclidine
(PBS): phosphate buffer saline
(PiVR): Raspberry Pi virtual reality system
(SSRI): selective serotonin reuptake inhibitor
(5-HT): serotonin
(SERT): serotonin transporter
(SEM): standard error of the mean
(t_50_): time to half max decay
(TRD): treatment resistant depression
(VNC): ventral nerve cord
(WT): wild-type
(8-OH-DPAT): 8-hydroxy-2-(di-n-propylamino) tetralin hydrobromide

## Acknowledgments

We thank Dr. Mimi Shin (University of Virginia, Chemistry department) and Dr. Jay Hirsh (Biology department) for their advice on feeding and locomotion assays. Dr. Jeffery Copeland (Eastern Mennonite University, Biology Department) provided guidance on fly crosses. We also thank Dr. Bita Moghaddam (Oregon Health & Science University, Behavioral Neuroscience Program) for advice on additional pharmacology comparison assays with ketamine. Dr. David Tadres (University of California, Santa Barbara, Biology Department) helped troubleshoot the PiVR Instrument. UVA’s MAE Rapid Prototyping and Machine lab was used for 3D printing PiVR parts. Cartoon figures were created in BioRender. This work was funded by NIH R01MH085159.

## Author contributions statement

K.E.D. and B.J.V. designed the study, analyzed and interpreted data, and wrote and edited the manuscript. K.E.D. conducted most of the experiments. K.H.K and L.W. assisted with feeding and locomotion assays and optogenetic experiments. B.J.V. secured funding.

## Supporting Information

### I. Supplemental Figures

**Figure S1.**
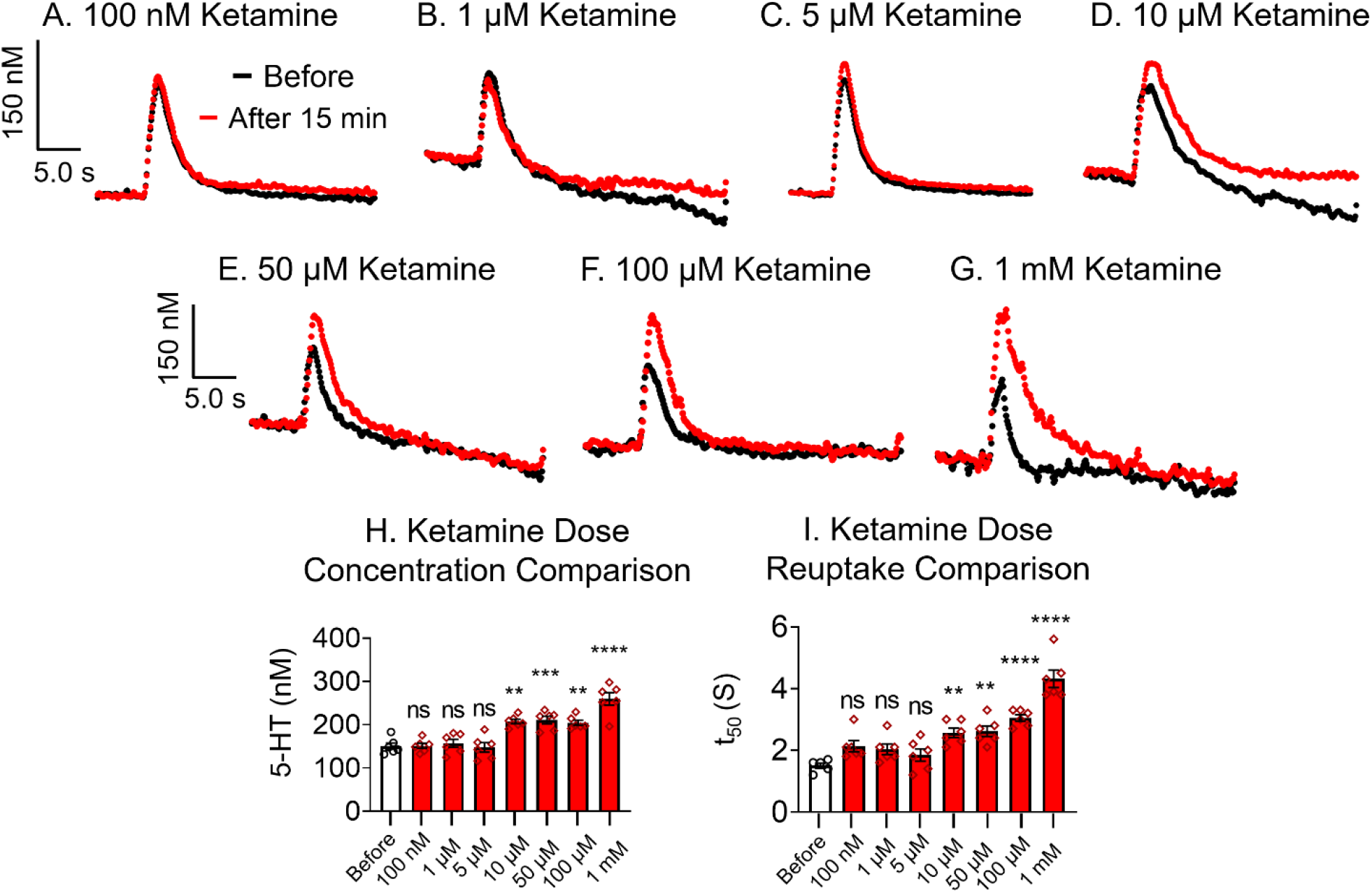
Serotonin release and reuptake effects for acute ketamine (*n* = 6 larvae each). Ketamine was bath-applied to *Drosophila* larvae VNC tissue and optogenetic stimulation repeated every 5 mins for 15 mins. Serotonin concentration and reuptake are characterized by conc. versus *t* plots before (black) and after (**A**) 100 nM, (**B**) 1 µM, (**C**) 5 µM, (**D**) 10 µM, (**E**) 50 µM, (**F**) 100 µM, and (**G**) 1 mM ketamine. (**H**) Evoked concentration of serotonin vs ketamine dose. There was a significant overall effect of ketamine dose on serotonin concentration (One-Way ANOVA, F_(7,40)_ = 20.51, *p* ≤ 0.0001, *n* = 6), and Tukey’s post-hoc determined serotonin concentrations were significantly different than pre-drug at 10 µM (p ˂ 0.0014), 50 µM (p ˂ 0.0007), 100 µM (p ˂ 0.0029), and 1 mM ketamine (*p* ≤ 0.0001). (**I**) For serotonin reuptake, there was a significant overall effect of ketamine dose on serotonin reuptake (One-Way ANOVA F_(7,40)_ = 24.48, *p* ≤ 0.0001, *n* = 6). Tukey’s post-hoc revealed reuptake increased significantly with 10 µM (p ˂ 0.0058), 50 µM (p ˂ 0.0034), 100 µM (*p* ˂ 0.0001), and 1 mM ketamine (*p* ˂ 0.0001).

**Figure S2.**
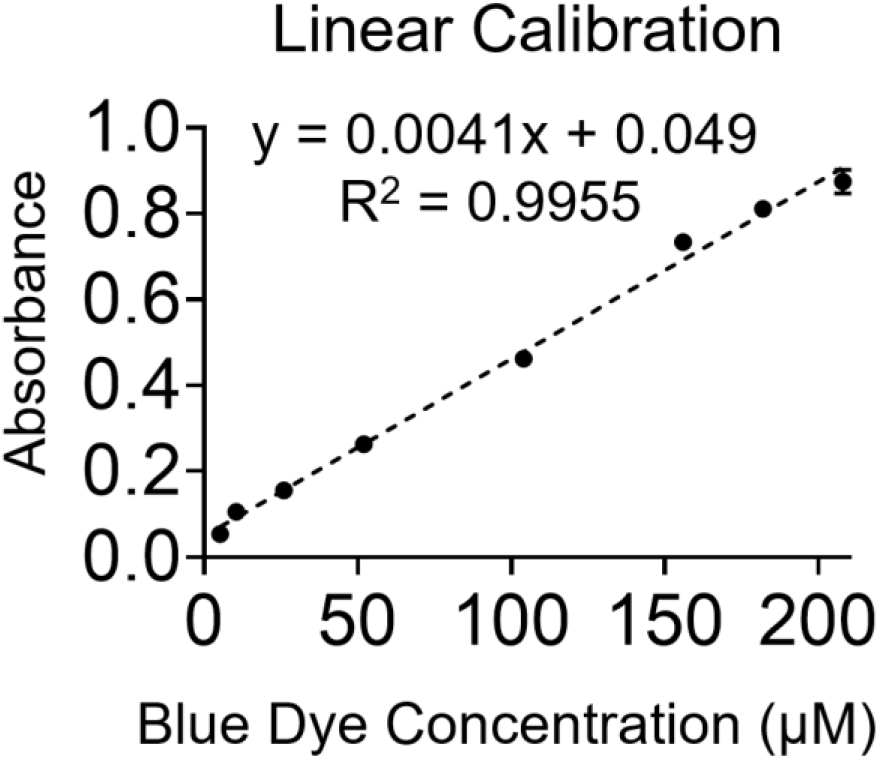
Blue dye linear calibration for *Drosophila* larvae food consumption (*n* = 3 replicates/ concentration). The linear equation was used to back-calculate the mass (µg) of food eaten per larva per day.

**Figure S3.**
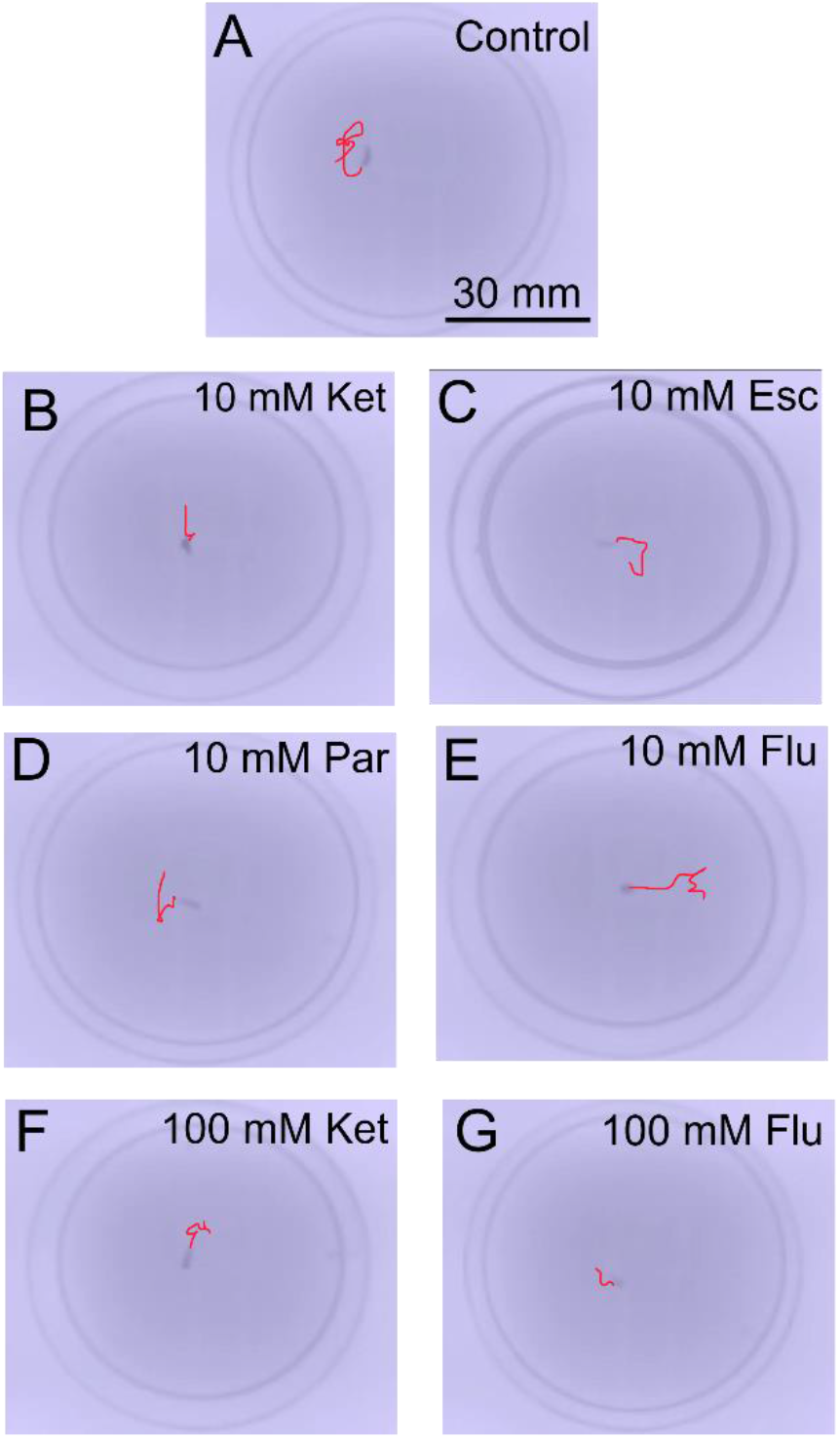
Locomotion tracking comparison after *Drosophila* larvae ate different antidepressant drugs for 24 hours (*n* = 30 larvae/dose). A larva’s locomotion was tracked for 60 s after (A) PBS, (B) 10 mM ketamine, (C) 10 mM escitalopram, (D) 10 mM paroxetine, (E) 10 mM fluoxetine, (F) 100 mM ketamine, and (G) 100 mM fluoxetine. One-Way ANOVA (F_(10,319)_ = 72.35, *p* ≤ 0.0001, *n* = 30) and Tukey’s post-hoc comparisons found the distance traveled significantly increased with 10 mM fluoxetine (p****< 0.0001). However, distance significantly decreased with 100 mM fluoxetine (p ≥ 0.0020).

**Figure S4.**
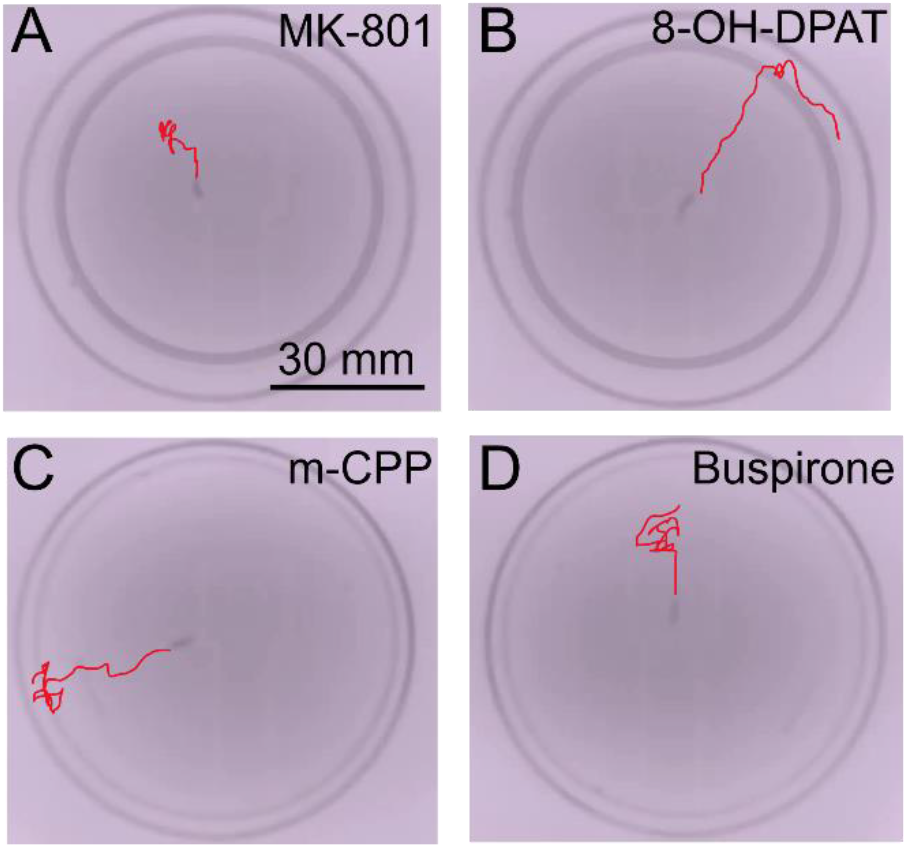
Locomotion tracking comparison after *Drosophila* larvae ate different drugs for 24 hours compared to ketamine and SSRIs (*n* = 30 larvae/dose). A larva’s locomotion was tracked for 60 s after (**A**) 1 mM MK-801, (**B**) 1 mM 8-OH-DPAT, (**C**) 1 mM m-CPP, and (**D**) 1 mM buspirone. One-Way ANOVA (F_(8,261)_ = 80.72, *p* ≤ 0.0001, *n* = 30) and Tukey’s post-hoc comparisons found the distance traveled significantly increased the most with 8-OH-DPAT, m-CPP, and buspirone (all p****< 0.0001) and slightly increased with MK-801 (p*< 0.1). Compared to 1 mM ketamine, locomotion was not different with m-CPP or buspirone (both p˃ 0.1736). However, larvae moved more with 1 mM 8-OH-DPAT (p*< 0.1) compared to ketamine. With 1 mM MK-801, distance traveled was less than with 1 mM ketamine (p****< 0.0001).

## References

Barann, M., Stamer, U. M., Lyutenska, M., Stüber, F., Bönisch, H., & Urban, B. (2014). Effects of opioids on human serotonin transporters. Naunyn-Schmiedeberg’s Archives of Pharmacology, 388(1), 43–49. 10.1007/s00210-014-1056-3

Berman, R. M., Cappiello, A., Anand, A., Oren, D. A., Heninger, G. R., Charney, D. S., & Krystal, J. H. (2000). Antidepressant effects of ketamine in depressed patients. Biological Psychiatry, 47(4), 351–354. 10.1016/S0006-3223(99)00230-9

Borue, X., Condron, B., & Venton, B. J. (2010). Both synthesis and reuptake are critical for replenishing the releasable serotonin pool in Drosophila. Journal of Neurochemistry, 113(1), 188–199. 10.1111/j.1471-4159.2010.06588.x

Borue, X., Cooper, S., Hirsh, J., Condron, B., & Venton, B. J. (2009). Quantitative evaluation of serotonin release and clearance in Drosophila. Journal of Neuroscience Methods, 179(2), Article 2. 10.1016/j.jneumeth.2009.02.013

Bowman, M. A., Vitela, M., Clarke, K. M., Koek, W., & Daws, L. C. (2020). Serotonin transporter and plasma membrane monoamine transporter are necessary for the antidepressant-like effects of ketamine in mice. International Journal of Molecular Sciences, 21(20), 1–22. 10.3390/ijms21207581

Can, A., Zanos, P., Moaddel, R., Kang, H. J., Dossou, K. S. S., Wainer, I. W., Cheer, J. F., Frost, D. O., Huang, X. P., & Gould, T. D. (2016). Effects of ketamine and ketamine metabolites on evoked striatal dopamine release, dopamine receptors, and monoamine transporters. Journal of Pharmacology and Experimental Therapeutics, 359(1), 159–170. 10.1124/jpet.116.235838

De Boer, S. F., & Koolhaas, J. M. (2005). 5-HT1A and 5-HT1B receptor agonists and aggression: A pharmacological challenge of the serotonin deficiency hypothesis. European Journal of Pharmacology, 526(1–3), 125–139. 10.1016/j.ejphar.2005.09.065

De Vry, J., & Schreiber, R. (2000). Effects of selected serotonin 5-HT1 and 5-HT2 receptor agonists on feeding behavior: Possible mechanisms of action. Neuroscience and Biobehavioral Reviews, 24(3), 341–353. 10.1016/S0149-7634(99)00083-4

Dumitrescu, E., Copeland, J. M., & Venton, B. J. (2023). *Parkin* Knockdown Modulates Dopamine Release in the Central Complex, but Not the Mushroom Body Heel, of Aging *Drosophila*. ACS Chemical Neuroscience, 14(2), 198–208. 10.1021/acschemneuro.2c00277

Dunham, K. E., & Venton, B. J. (2020). Improving serotonin fast-scan cyclic voltammetry detection: New waveforms to reduce electrode fouling. Analyst, 145(22), 7437–7446. 10.1039/d0an01406k

Dunham, K. E., & Venton, B. J. (2022). SSRI antidepressants differentially modulate serotonin reuptake and release in Drosophila. *Journal of Neurochemistry*, June, 1–13. 10.1111/jnc.15658

El Mansari, M., Sánchez, C., Chouvet, G., Renaud, B., & Haddjeri, N. (2005). Effects of Acute and Long-Term Administration of Escitalopram and Citalopram on Serotonin Neurotransmission: An In Vivo Electrophysiological Study in Rat Brain. Neuropsychopharmacology, 30(7), 1269–1277. 10.1038/sj.npp.1300686

Fakhoury, M. (2016). Revisiting the Serotonin Hypothesis: Implications for Major Depressive Disorders. Molecular Neurobiology, 53(5), 2778–2786. 10.1007/s12035-015-9152-z

Galvanho, J. P., Manhães, A. C., Carvalho-Nogueira, A. C. C., Silva, J. de M., Filgueiras, C. C., & Abreu-Villaça, Y. (2020). Profiling of behavioral effects evoked by ketamine and the role of 5HT2 and D2 receptors in ketamine-induced locomotor sensitization in mice. Progress in Neuro-Psychopharmacology and Biological Psychiatry, 97, 109775. 10.1016/j.pnpbp.2019.109775

Gould, G. G., Hensler, J. G., Burke, T. F., Benno, R. H., Onaivi, E. S., & Daws, L. C. (2011). Density and function of central serotonin (5-HT) transporters, 5-HT1A and 5-HT2A receptors, and effects of their targeting on BTBR T+tf/J mouse social behavior. Journal of Neurochemistry, 116(2), 291–303. 10.1111/j.1471-4159.2010.07104.x

Hales, K. G., Korey, C. A., Larracuente, A. M., & Roberts, D. M. (2015). Genetics on the fly: A primer on the drosophila model system. Genetics, 201(3), 815–842. 10.1534/genetics.115.183392

Hashemi, P., Dankoski, E. C., Lama, R., Wood, K. M., Takmakov, P., & Wightman, R. M. (2012). Brain dopamine and serotonin differ in regulation and its consequences. Proceedings of the National Academy of Sciences of the United States of America, 109(29), 11510–11515. 10.1073/pnas.1201547109

Hashemi, P., Dankoski, E. C., Petrovic, J., Keithley, R. B., & Wightman, R. M. (2009). Voltammetric detection of 5 -hydroxytryptamine release in the rat brain. Analytical Chemistry, 81(22), 9462–9471. 10.1021/ac9018846

Hidalgo, S., Molina-Mateo, D., Escobedo, P., Zárate, R. V., Fritz, E., Fierro, A., Perez, E. G., Iturriaga-Vasquez, P., Reyes-Parada, M., Varas, R., Fuenzalida-Uribe, N., & Campusano, J. M. (2017). Characterization of a Novel Drosophila SERT Mutant: Insights on the Contribution of the Serotonin Neural System to Behaviors. ACS Chemical Neuroscience, 8(10), 2168–2179. 10.1021/acschemneuro.7b00089

Irifune, M., Shimizu, T., & Nomoto, M. (1991). Ketamine-induced hyperlocomotion associated with alteration of presynaptic components of dopamine neurons in the nucleus accumbens of mice. Pharmacology Biochemistry and Behavior, 40(2), 399–407. 10.1016/0091-3057(91)90571-I

Jeibmann, A., & Paulus, W. (2009). Drosophila melanogaster as a model organism of brain diseases. International Journal of Molecular Sciences, 10(2), 407–440. 10.3390/ijms10020407

Johnson, O., Becnel, J., & Nichols, C. D. (2009). Serotonin 5-HT2 and 5-HT1A-like receptors differentially modulate aggressive behaviors in Drosophila melanogaster. Neuroscience, 158(4), 1292–1300. 10.1016/j.neuroscience.2008.10.055

Kasture, A. S., Hummel, T., Sucic, S., & Freissmuth, M. (2018). Big lessons from tiny flies: Drosophila melanogaster as a model to explore dysfunction of dopaminergic and serotonergic neurotransmitter systems. International Journal of Molecular Sciences, 19(6). 10.3390/ijms19061788

Keszthelyi, D., Troost, F. J., & Masclee, A. A. M. (2009). Understanding the role of tryptophan and serotonin metabolism in gastrointestinal function. Neurogastroenterology and Motility, 21(12), 1239–1249. 10.1111/j.1365-2982.2009.01370.x

Knapp, E. M., Kaiser, A., Arnold, R. C., Sampson, M. M., Ruppert, M., Xu, L., Anderson, M. I., Bonanno, S. L., Scholz, H., Donlea, J. M., & Krantz, D. E. (2022). Mutation of the Drosophila melanogaster serotonin transporter dSERT impacts sleep, courtship, and feeding behaviors. PLOS Genetics, 18(11), e1010289. 10.1371/journal.pgen.1010289

Koksal, P. M., & Gürbüzel, M. (2015). Analysis of genotoxic activity of ketamine and rocuronium bromide using the somatic mutation and recombination test in Drosophila melanogaster. Environmental Toxicology and Pharmacology, 39(2), 628–634. 10.1016/j.etap.2014.12.010

Kraus, C., Wasserman, D., Henter, I. D., Acevedo-Diaz, E., Kadriu, B., & Zarate, C. A. (2019). The influence of ketamine on drug discovery in depression. Drug Discovery Today, 24(10), 2033–2043. 10.1016/j.drudis.2019.07.007

Lawal, H. O., Terrell, A., Lam, H. A., Djapri, C., Jang, J., Hadi, R., Roberts, L., Shahi, V., Chou, M. T., Biedermann, T., Huang, B., Lawless, G. M., Maidment, N. T., & Krantz, D. E. (2014). Drosophila modifier screens to identify novel neuropsychiatric drugs including aminergic agents for the possible treatment of Parkinson’s disease and depression. Molecular Psychiatry, 19(2), 235–242. 10.1038/mp.2012.170

Li, Y. F. (2020). A hypothesis of monoamine (5-HT) – Glutamate/GABA long neural circuit: Aiming for fast-onset antidepressant discovery. Pharmacology and Therapeutics, 208, 107494. 10.1016/j.pharmthera.2020.107494

Liu, W. W., & Wilson, R. I. (2013). Glutamate is an inhibitory neurotransmitter in the *Drosophila* olfactory system. Proceedings of the National Academy of Sciences, 110(25), 10294–10299. 10.1073/pnas.1220560110

Majeed, Z. R., Abdeljaber, E., Soveland, R., Cornwell, K., Bankemper, A., Koch, F., & Cooper, R. L. (2016). Modulatory Action by the Serotonergic System: Behavior and Neurophysiology in Drosophila melanogaster. Neural Plasticity, 2016. 10.1155/2016/7291438

Martin, C. A., & Krantz, D. E. (2014). Drosophila melanogaster as a genetic model system to study neurotransmitter transporters. Neurochemistry International, 73(1), 71–88. 10.1016/j.neuint.2014.03.015

Marvin, J. S., Scholl, B., Wilson, D. E., Podgorski, K., Kazemipour, A., Muller, J. A., Schoch, S., Quiroz, F. J. U., Rebola, N., Bao, H., Little, J. P., Tkachuk, A. N., Cai, E., Hantman, A. W., Wang, S. S.-H., DePiero, V. J., Borghuis, B. G., Chapman, E. R., Dietrich, D., … Looger, L. L. (2018). Stability, affinity, and chromatic variants of the glutamate sensor iGluSnFR. Nature Methods, 15(11), Article 11. 10.1038/s41592-018-0171-3

Moghaddam, B., Adams, B., Verma, A., & Daly, D. (1997). Activation of Glutamatergic Neurotransmission by Ketamine: A Novel Step in the Pathway from NMDA Receptor Blockade to Dopaminergic and Cognitive Disruptions Associated with the Prefrontal Cortex. The Journal of Neuroscience, 17(8), 2921–2927. 10.1523/JNEUROSCI.17-08-02921.1997

Ni, Y. G., & Miledi, R. (1997). Blockage of 5HT _2C_ serotonin receptors by fluoxetine (Prozac). Proceedings of the National Academy of Sciences, 94(5), 2036–2040. 10.1073/pnas.94.5.2036

Owens, M. J., Knight, D. L., & Nemeroff, C. B. (2001). Second-generation SSRIs: Human monoamine transporter binding profile of escitalopram and R-fluoxetine. Biological Psychiatry, 50(5), 345–350. 10.1016/S0006-3223(01)01145-3

Pham, T. H., Mendez-David, I., Defaix, C., Guiard, B. P., Tritschler, L., David, D. J., & Gardier, A. M. (2017). Ketamine treatment involves medial prefrontal cortex serotonin to induce a rapid antidepressant -like activity in BALB/cJ mice. Neuropharmacology, 112, 198–209. 10.1016/j.neuropharm.2016.05.010

Pörzgen, P., Park, S. K., Hirsh, J., Sonders, M. S., & Amara, S. G. (2001). The antidepressant-sensitive dopamine transporter in Drosophila melanogaster: A primordial carrier for catecholamines. Molecular Pharmacology, 59(1), 83–95. 10.1124/mol.59.1.83

Privman, E., & Venton, B. J. (2015). Comparison of dopamine kinetics in the larval Drosophila ventral nerve cord and protocerebrum with improved optogenetic stimulation. Journal of Neurochemistry, 135(4), 695–704. 10.1111/jnc.13286

Puthongkham, P., Lee, S. T., & Venton, B. J. (2019). Mechanism of histamine oxidation and electropolymerization at carbon electrodes. Analytical Chemistry, 91(13), Article 13. 10.1021/acs.analchem.9b01178

Ragnhildstveit, A., Slayton, M., Jackson, L. K., Brendle, M., Ahuja, S., Holle, W., Moore, C., Sollars, K., Seli, P., & Robison, R. (2022). Ketamine as a Novel Psychopharmacotherapy for Eating Disorders: Evidence and Future Directions. Brain Sciences, 12(3). 10.3390/brainsci12030382

Ribaudo, G., Bortoli, M., Witt, C. E., Parke, B., Mena, S., Oselladore, E., Zagotto, G., Hashemi, P., & Orian, L. (2021). ROS-Scavenging Selenofluoxetine Derivatives Inhibit In Vivo Serotonin Reuptake. ACS Omega. 10.1021/acsomega.1c05567

Richter, F. G., Fendl, S., Haag, J., Drews, M. S., & Borst, A. (2018). Glutamate Signaling in the Fly Visual System. IScience, 7, 85–95. 10.1016/j.isci.2018.08.019

Ries, A. S., Hermanns, T., Poeck, B., & Strauss, R. (2017). Serotonin modulates a depression -like state in Drosophila responsive to lithium treatment. Nature Communications, 8, 1–11. 10.1038/ncomms15738

Roberts, D. B. (2006). Drosophila melanogaster: The model organism. Entomologia Experimentalis et Applicata, 121(2), 93–103. 10.1111/j.1570-8703.2006.00474.x

Robison, R., Lafrance, A., Brendle, M., Smith, M., Moore, C., Ahuja, S., Richards, S., Hawkins, N., & Strahan, E. (2022). A case series of group-based ketamine-assisted psychotherapy for patients in residential treatment for eating disorders with comorbid depression and anxiety disorders. Journal of Eating Disorders, 10(1), 1–9. 10.1186/s40337-022-00588-9

Ruberto, V. L., Jha, M. K., & Murrough, J. W. (2020). Pharmacological treatments for patients with TRD. Pharmaceuticals, 13, 116.

Saylor, R. A., Hersey, M., West, A., Buchanan, A. M., Berger, S. N., Nijhout, H. F., Reed, M. C., Best, J., & Hashemi, P. (2019). In vivo hippocampal serotonin dynamics in male and female mice: Determining effects of acute escitalopram using fast scan cyclic voltammetry. Frontiers in Neuroscience, 13(APR), 1–13. 10.3389/fnins.2019.00362

Schwartz, T., Trunko, M. E., Feifel, D., Lopez, E., Peterson, D., Frank, G. K. W., & Kaye, W. (2021). A longitudinal case series of IM ketamine for patients with severe and enduring eating disorders and comorbid treatment-resistant depression. Clinical Case Reports, 9(5), 1–7. 10.1002/ccr3.3869

Shell, B. C., & Grotewiel, M. (2022). Identification of additional dye tracers for measuring solid food intake and food preference via consumption-excretion in Drosophila. Scientific Reports, 12(1), 1–12. 10.1038/s41598-022-10252-6

Shell, B. C., Luo, Y., Pletcher, S., & Grotewiel, M. (2021). Expansion and application of dye tracers for measuring solid food intake and food preference in Drosophila. Scientific Reports, 11(1), 1–13. 10.1038/s41598-021-99483-7

Shell, B. C., Schmitt, R. E., Lee, K. M., Johnson, J. C., Chung, B. Y., Pletcher, S. D., & Grotewiel, M. (2018). Measurement of solid food intake in Drosophila via consumption-excretion of a dye tracer. Scientific Reports, 8(1), 1–13. 10.1038/s41598-018-29813-9

Shin, M., Copeland, J. M., & Venton, B. J. (2020). Real-Time Measurement of Stimulated Dopamine Release in Compartments of the Adult Drosophila melanogaster Mushroom Body. BioRxiv, 2020.06.29.177675. 10.1101/2020.06.29.177675

Silva, B., Goles, N. I., Varas, R., & Campusano, J. M. (2014). Serotonin receptors expressed in Drosophila mushroom bodies differentially modulate larval locomotion. PLoS ONE, 9(2). 10.1371/journal.pone.0089641

Spies, M., James, G. M., Berroterán-Infante, N., Ibeschitz, H., Kranz, G. S., Unterholzner, J., Godbersen, M., Gryglewski, G., Hienert, M., Jungwirth, J., Pichler, V., Reiter, B., Silberbauer, L., Winkler, D., Mitterhauser, M., Stimpfl, T., Hacker, M., Kasper, S., & Lanzenberger, R. (2018). Assessment of Ketamine binding of the serotonin transporter in humans with positron emission tomography. International Journal of Neuropsychopharmacology, 21(2), 145–153. 10.1093/ijnp/pyx085

Stahl, S. M. (1998). Mechanism of action of serotonin selective reuptake inhibitors. Journal of Affective Disorders, 51(3), 215–235. 10.1016/S0165-0327(98)00221-3

Stahl, S. M. (2013). Mechanism of action of ketamine. CNS Spectrums, 18(4), 171–174. 10.1017/S109285291300045X

Tadres, D., & Louis, M. (2020). PiVR: An affordable and versatile closed-loop platform to study unrestrained sensorimotor behavior. PLoS Biology, 18(7), 1–25. 10.1371/journal.pbio.3000712

Verma, A., & Moghaddam, B. (n.d.). NMDA Receptor Antagonists Impair Prefrontal Cortex Function as Assessed via Spatial Delayed Alternation Performance in Rats: Modulation by bopamine *-*.

West, A. M., Holleran, K. M., & Jones, S. R. (2023). Kappa Opioid Receptors Reduce Serotonin Uptake and Escitalopram Efficacy in the Mouse Substantia Nigra Pars Reticulata. International Journal of Molecular Sciences, 24(3), 2080. 10.3390/ijms24032080

Wood, K. M., & Hashemi, P. (2013). Fast-scan cyclic voltammetry analysis of dynamic serotonin reponses to acute escitalopram. ACS Chemical Neuroscience, 4(5), Article 5. 10.1021/cn4000378

Wood, K. M., Zeqja, A., Nijhout, H. F., Reed, M. C., Best, J., & Hashemi, P. (2014). Voltammetric and mathematical evidence for dual transport mediation of serotonin clearance in vivo. Journal of Neurochemistry, 130(3), 351–359. 10.1111/jnc.12733

Zanos, P., Thompson, S. M., Duman, R. S., Zarate, C. A., & Gould, T. D. (2018). Convergent Mechanisms Underlying Rapid Antidepressant Action. CNS Drugs, 32(3), 197–227. 10.1007/s40263-018-0492-x

Zhong, H., Sánchez, C., & Caron, M. G. (2012). Consideration of allosterism and interacting proteins in the physiological functions of the serotonin transporter. Biochemical Pharmacology, 83(4), 435–442. 10.1016/j.bcp.2011.09.020

Zimmerman, J. E., Chan, M. T., Lenz, O. T., Keenan, B. T., Maislin, G., & Pack, A. I. (2017). Glutamate Is a Wake-Active Neurotransmitter in Drosophila melanogaster. Sleep, 40(2). 10.1093/sleep/zsw046

